# “A Fine-Tuned Phosphatidylinositol Profile Contributes to Colonocyte Differentiation and Malignization: Evidence From Integrated Omics”

**DOI:** 10.64898/2026.02.20.706976

**Authors:** Albert Maimó-Barceló, Joan Bestard-Escalas, Karim Pérez-Romero, Lucía Martín-Saiz, Josep Muncunill-Fortuny, Catalina Crespí, Marco A. Martínez, M. Laura Martin, Daniel H. Lopez, Gonzalo P. Martín, José M. Olea, José A. Fernández, Ramon M. Rodriguez, Gwendolyn Barceló-Coblijn

## Abstract

Membrane lipid composition changes concomitantly with human colonocyte differentiation, a tightly regulated process occurring along the colon crypt. This process is heavily disrupted in colon cancer. Nonetheless, the regulatory mechanisms driving these changes, especially the replacement of arachidonic acid phosphatidylinositol species with monounsaturated fatty acid species, and how they are altered in cancer, remain unknown. To establish the transcriptional networks underlying this remodeling, we integrated transcriptomic and lipidomic profiles of isolated healthy and tumor human colonocytes using system biology approaches; identifying key gene regulatory networks involved in arachidonic acid and eicosanoid metabolism and phosphatidylinositol cycle as significant regulators during differentiation. Consistently, a distinct impact was found on organoid differentiation depending on colonocyte subtype and specific prostaglandin. Remarkably, the shift and associated transcriptomic programs were lost in tumor that heightened phosphoinositide metabolism. Altogether, these results underscore the importance of lipid remodeling in colonocyte stemness maintenance and proper onset of differentiation programs.

## BACKGROUND

Colon physiology is highly dependent on the renewal capacity of the epithelium, relying upon colonocyte stem cells (CSC) located at the bottom of the crypt. CSC give rise to several lineages of mature colonocytes through a process that occurs gradually along the crypt, ending up at the apical side with death by anoikis(1,2) (**Figure 1a**). The colon epithelium differentiation involves several crucial signaling pathways, such as Wnt/β-catenin and BMP/TGF-β, which play vital roles in regulating CSC proliferation, differentiation, and maintenance within the crypts.(3,4) The tight regulation of differentiation is significantly impaired in colon cancer, which, together with the accumulation of molecular alterations, promotes the emergence and subsequent advancement of colon cancer-initiating cells.(5–7) These pathways have been studied in detail given the high incidence and death rate of colon cancer,(8) providing the evidence to develop novel and targeted therapies.(9) However, these studies disentangle the molecular insights at the protein and gene level, leaving the potential implication of metabolites, including lipids, unexplored.

**Figure 1.**
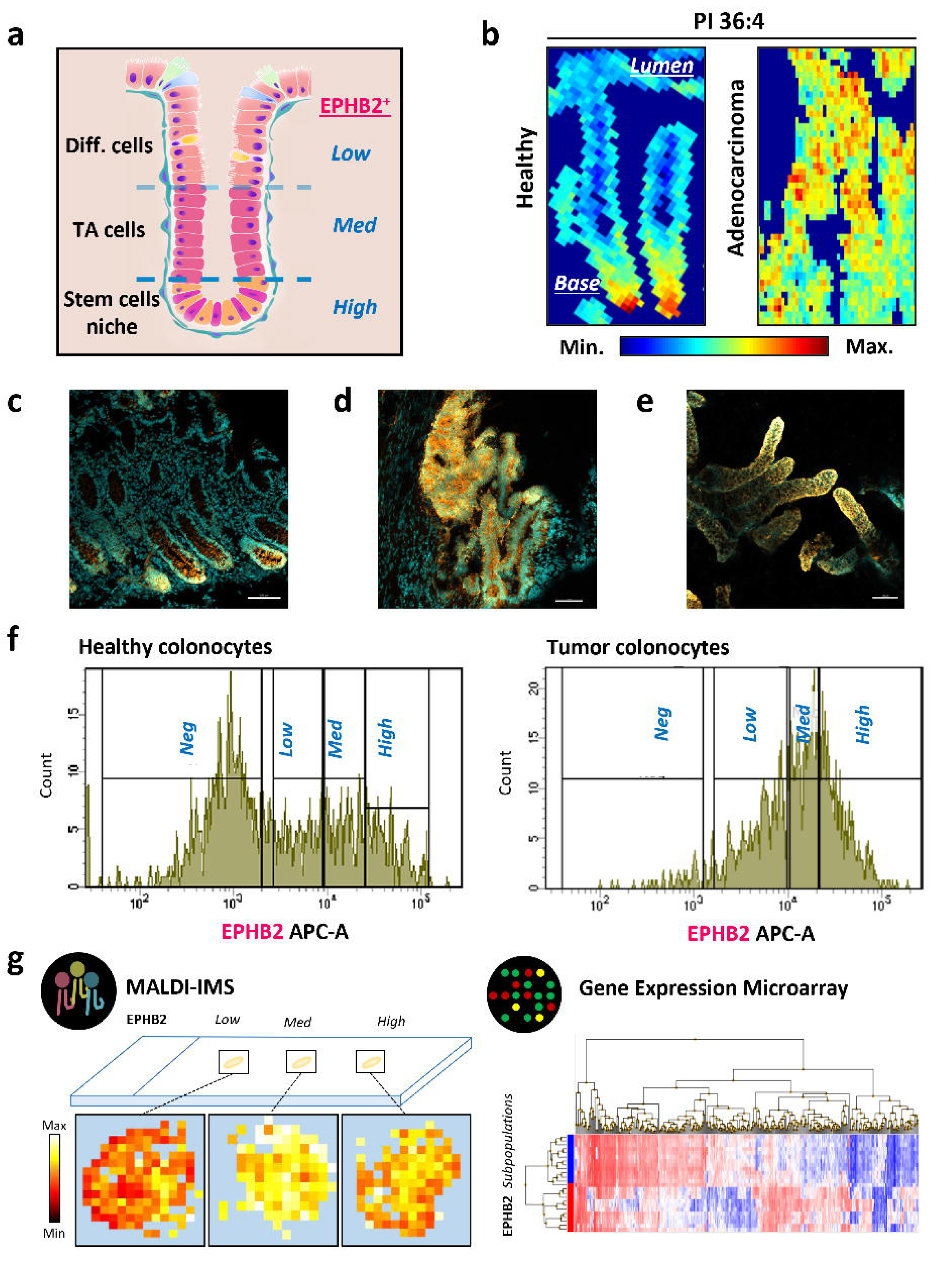
Lipid fingerprint faithfully conveys the physiological state of the colonocyte. a) Scheme summarizing the differentiation process along the colon crypt. b) Representative MALDI-MSI image showing the distribution of an AA-containing lipid species, PI 36:4, in human healthy and adenocarcinoma colon samples. c-d) Representative immunofluorescence (IF) images of histological sections showing EPHB2 gradual expression (gold) along healthy colon crypt (c) while in tumor (d), the pattern is compromised. e) IF of isolated crypts showing EPHB2 polarized distribution (DAPI, cyan). f) FACS isolated EPHB2 subpopulations: High, Med (medium), Low, and Neg (negative) EPHB2. g) Each subpopulation was analyzed either by MALDI-MSI or gene expression microarray. TA cells (Transit-amplifying cells). Scale bar = 100 µm.

**Figure 2.**
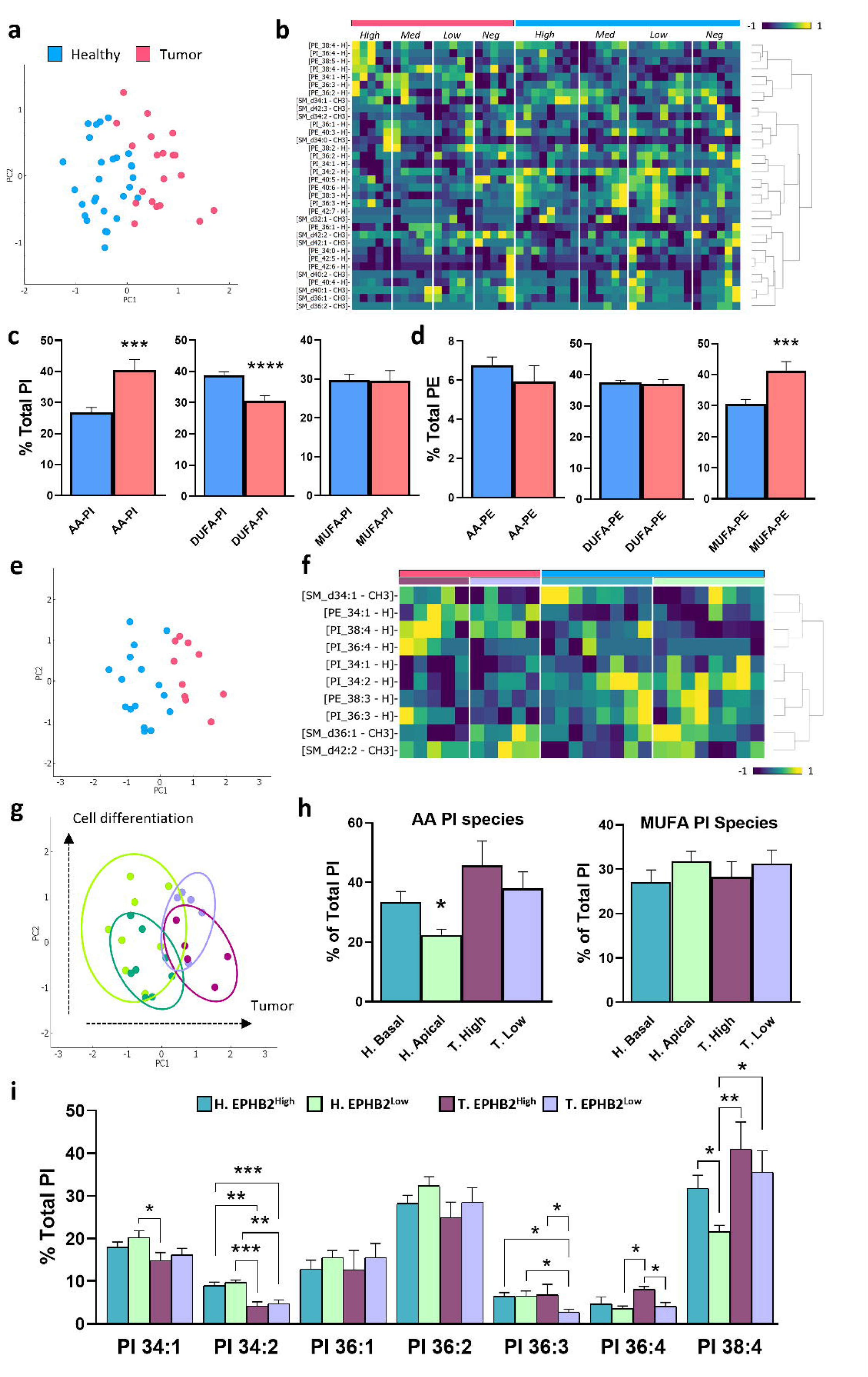
Differences in lipid species content in healthy and tumor colonocyte subpopulations. a) PCA and b) Hierarchical-clustering heatmap of lipid species comparing all healthy (blue) vs. all tumor (tumor) EPHB2 subpopulations. c) Bar plots comparing PI species levels in healthy *vs*. tumor colonocyte subpopulations. d) Bar plots comparing PE species levels in healthy *vs*. tumor colonocyte subpopulations. e) PCA plot of PI species composition in Low and High EPHB2 colonocyte subpopulations according to it pathological condition and f) Hierarchical-Clustering Heatmap of the top-10 ANOVA-ranked lipid species comparing healthy and tumor, both EPHB2^High^ and EPHB2^Low^; g) PCA plot showing differences between EPHB2^High^ and EPHB2^Low^ according to tumor origin and cell-differentiation-state; h) Bar plots comparing AA-PI and MUFA-PI species levels in healthy or tumor EPHB2^High^ vs. EPHB2^Low^ subpopulations (two-tailed t-test, \**p* <0.05). i) Bar plots comparing PI species levels in EPHB2^High^ vs. EPHB2^Low^ subpopulations in healthy (H) and tumor (T) samples. a-f) blue dots and bars, healthy samples; red, tumor samples. In bar diagrams, values are expressed as a percentage of total lipid class and represent the mean ± SEM (n=5-8). Statistical significance was assessed using multiple comparisons ordinary one-way ANOVA, \**p* <.05, \*\**p* <.01, \*\*\**p* <.001. AA: arachidonic acid; MUFA: monounsaturated fatty acids; PI: phosphatidylinositol; PE: phosphatidylethanolamine; SM: sphingomyelin. Extended lipidomic data in Supplementary Table 3.

Using spatially resolved imaging lipidomic techniques, we have described tissue- and cell-type-specific differences in lipid composition within and between healthy and tumor colon mucosa, unveiling unexpected layers of regulation at the composition level(10). We uncovered that the gradual distribution of arachidonic acid (AA)- and monounsaturated fatty acid (MUFA)-containing phospholipids along the healthy basal-apical axis of the mucosa adjusts to simple mathematical equations, and changes concomitantly with colonocyte differentiation(11,12) (**Figure 1b, Supplementary Table 1**). Further, the phospholipid fingerprint is sustained by the gradual expression of key lipid enzymes, in particular those accounting for AA handling.(11–13) Consistent with this strict regulation in healthy epithelium, lipid and protein gradients are lost in tumor.(11,12,14,15) Altogether, these results point to a tight relationship between the specific colonocyte lipid profile and both differentiation and malignization states, underscoring the importance of membrane lipid composition in cell homeostasis. However, knowledge regarding how this specificity is achieved, that is, the signaling pathways and mechanisms acting at the genetic and enzymatic regulatory level, is scant.(16–18)

The study of regulatory mechanisms at the lipid species level is highly complex because of the specific features of lipids as biomolecules and their metabolism(17). Thus, the enzymatic networks controlling the level of each one of the lipid species present in the cell are complex and remain poorly characterized. Moreover, many of these enzymes are both redundant and promiscuous, complicating the dissection of lipid metabolism because perturbation of a single lipid enzyme can affect simultaneously numerous lipid species. Considering that at least 26 phospholipid species were previously identified to change gradually along the crypt, we decided to take a multi-omic data integration approach to delve into the regulation of the MUFA-/AA-lipid species gradient (Figure 1). While the results offered a multifaceted scenario, it demonstrates the great potential of this approach to generate novel hypotheses addressing complex pathophysiological processes.

## METHODS

### Tissue histological sections and immunofluorescence

Sections of approx.10 μm thickness were prepared using no cryoprotective and no embedding substances in a cryostat (Leica CM3050S, Leica Biosystems, Germany) at −20 °C. Tissue sections were fixed with ice-cold methanol:acetone (1:1, v, v) (Sigma-Aldrich, Spain), washed with phosphate-buffered saline (PBS), followed by incubation with 0.5% BSA (Sigma-Aldrich, Spain) in PBS for 40 min at room temperature (RT). Immunostaining with monoclonal EPHB2-APC mouse anti-human (0.1 tests/section) (2H9 clone, BD biosciences, Spain) diluted 1:100 in 0.2% BSA-PBS was carried out for 1 h at room temperature in a humidified chamber. Samples were washed several times with TBS 0.01 % Tween (Sigma-Aldrich, Spain). Nuclei staining was performed using Fluoroshield™ with 4′,6-diamidino-2-phenylindole (DAPI) (Sigma-Aldrich, Spain) as histology mounting medium. Finally, samples were observed with LSM 700 confocal microscope (Carl Zeiss, Germany).

### Isolation of human colon crypts

The protocol was adapted from Merlos-Suárez *et al*.(7) Briefly, healthy and tumor colon crypts were isolated from surgical biopsies of patients with CC **(Supplementary table 2)**. Samples were washed with standard PBS, cut into 1-2 cm fragments, and incubated with 10 μM DTT (Sigma-Aldrich, Spain) for 5 min at RT, and with 8 mM EDTA-HBSS (Sigma-Aldrich, Spain) for 45-60 min at 4 °C. Supernatant was replaced with 10 ml of fresh 5% FBS-HBSS (Sigma-Aldrich, Spain), and crypts were isolated by a 4-min vigorously shaking of the colon fractions at least 3 times. The crypt-enriched supernatant was centrifuged (100×g, 4 °C, 10 min) and washed three times.

### Immunofluorescence of freshly isolated crypts

Approximately·200 crypts were resuspended in 5% FBS-HBSS and incubated for 30 min at 4 °C with EPHB2 (2 test, APC Mouse Anti-Human, Clone 2H9, BD Biosciences, Barcelona, Spain). Crypts were washed 3 times with 200 μl PBS (100×g, 10 min, 4 °C), and 1 μl of DAPI Solution (1mg/ml, BD-Biosciences, Spain) was added at the end. Finally, 25 μl of the PBS containing the labeled crypts were placed on an adherent plain glass microscope slides positively charged and observed with confocal microscopy.

### Fluorescence-activated cell sorting (FACS)

Isolated crypts were disaggregated into single cell-colonocyte suspension following chemical digestion with TrypLE™ Express Enzyme (Gibco, A1285901) and DNAse I (200U/ml, 1:400, Roche, Germany) for 10-15 min at 37 °C, followed by further mechanical disaggregation using P1000 and P200 pipetting. Digestion was neutralized using FBS (5% final volume). Cell suspension was filtered through 100- and 40 μm cell strainer and single-cells recovered at 100×g, 10 min, 4 °C. Pellet was resuspended in 250 μl of SB and incubated 45 min at 4 °C with the following mix of antibodies: CD31 (2 test, PE-Cy™7, Mouse Anti-Human CD31, Clone WM59), CD45 (2 test, PE-Cy™7 mouse anti-human, Clone HI30), CD11b (2 test, PE-Cy™7 mouse anti-human, Clone ICRF44), CD117, EpCAM (2 test of 3 μg/ml, PE mouse anti-human, Clone EBA-1, RUO GMP), EPHB2 (2 test, APC Mouse Anti-Human, Clone 2H9) (BD Biosciences, Spain). Samples were washed 3 times and 1:300 DAPI Solution (1mg/ml, BD-Biosciences) was added before flow cytometry analysis. Cells showing EPCAM^+^, CD11b^−^, CD45^−^, CD31^−^, CD117^−^ staining were sorted into 4 different colonocyte subpopulations of approximately 1.5·10^4^ cells, according to EPHB2^+^ intensity (High, Med, Low and Neg EPHB2), reflecting colonocyte differentiation state **(Figure 1f, Supplementary Figure 12).**

### MALDI-MSI of lipids

The isolated EPHB2 subpopulations from healthy (n=8) and tumor samples (n=5) were analyzed by MALDI-MSI. Briefly, samples were thawed and immediately pelleted by centrifugation (300×g, 15 min, 4 °C). Cell pellets were resuspended by pipetting in 10 μl of PBS. Each sample was seeded on lysine-coated glass slides (0.2% lysine solution) by applying 0.5-1 μl drops, creating an overlapping spot of highly concentrated sample suitable for the analysis (**Figure 1g**, left).(19) The matrix 1,5-diaminonaphtalene (Sigma-Aldrich, Germany), was sublimated using the system developed by Fernandez et al.(20) Samples were analyzed at SGiker-UPV/EHU(Spain) in negative-ion-mode with MALDI-LTQ-Orbitrap XL analyzer (Thermo Fisher, San Jose, CA, USA) equipped with a MNL 100 N_2_ laser (LTB, Berlin), elliptical spot, 60 Hz repetition rate, optical arrangement of two mirrors and focusing lens of f=125 mm. Elliptical laser spot of 30–60 μJ, step-size of 50 μm. The scanning range was 400–1,200 Da and mass resolutions of 30,000 at m/z = 400 Da on FACS cells. Acquisition of MALDI-MSI data was performed as previously described in Garate et *al.*(10) Post-Imaging data analysis was carried using K-means or HD-RCA(10,21,22) from in house MatLAB^TM^ algorithms to define regions of interest. Peak intensity was normalized to the total ion current. Inter-experimental data were aligned as previously described in Maimó-Barceló et al.(23) and, lipid assignation was based on www.lipidmaps.org database, with a mass accuracy better than 9 ppm. Detailed information of colon epithelium LC–MS/MS and MALDI MS/MS can be found in previous publications(11,23,24). Assigned lipid molecular species abundances were normalized to the total lipid content within each respective lipid class, expressed as molecular species percentage of the class total. Further analysis of assigned data was performed using K-means analysis, PCA and hierarchical-clustering heatmap using Orange data mining (v.3.34). Statistical analysis and diagram plots were created with Prism 8 (Graphpad software, Inc). Data available in **Supplementary Table 3**.

### Gene expression data

The commercial gene expression microarray Human Clariom S pico (ThermoFisher Scientific, Spain) was used to characterize isolated EPHB2 subpopulations EPHB2 -Low, -med and -High) from patient-paired healthy and tumor samples (n=4) (**Figure 1g**, right), following the manufacturer instructions. RNA was extracted using RNeasy Micro Kit (QIAGEN, Spain), followed by assessments of RNA purity (260/280 ratio=1.7-2.0, Synergy H1, BioTek, Germany) and integrity (2100 Bioanalyzer system, Agilent Technologies, Spain), following the standard protocols. Microarray gene expression data was processed with Transcriptome analysis console (TAC, v.4) software following standard procedure. Gene Set Enrichment Analysis (GSEA) was performed using GSEA software v4.3.2 (**Figure 3**).(25) Software parameters were set as: chip collapsed, mean div norm, weighted score, 1,000 permutations gene set, t-test metric, and standard significance FDR <25% in all comparisons. Lipid-related gene list: Gene Ontology, KEGG, Reactome and Wikipathways databases were used to construct a detailed list of 1182 genes related to lipid processes and pathways **(Supplementary table 5).** Gene expression dataset is available at gene expression omnibus accession GSE285090.

**Figure 3.**
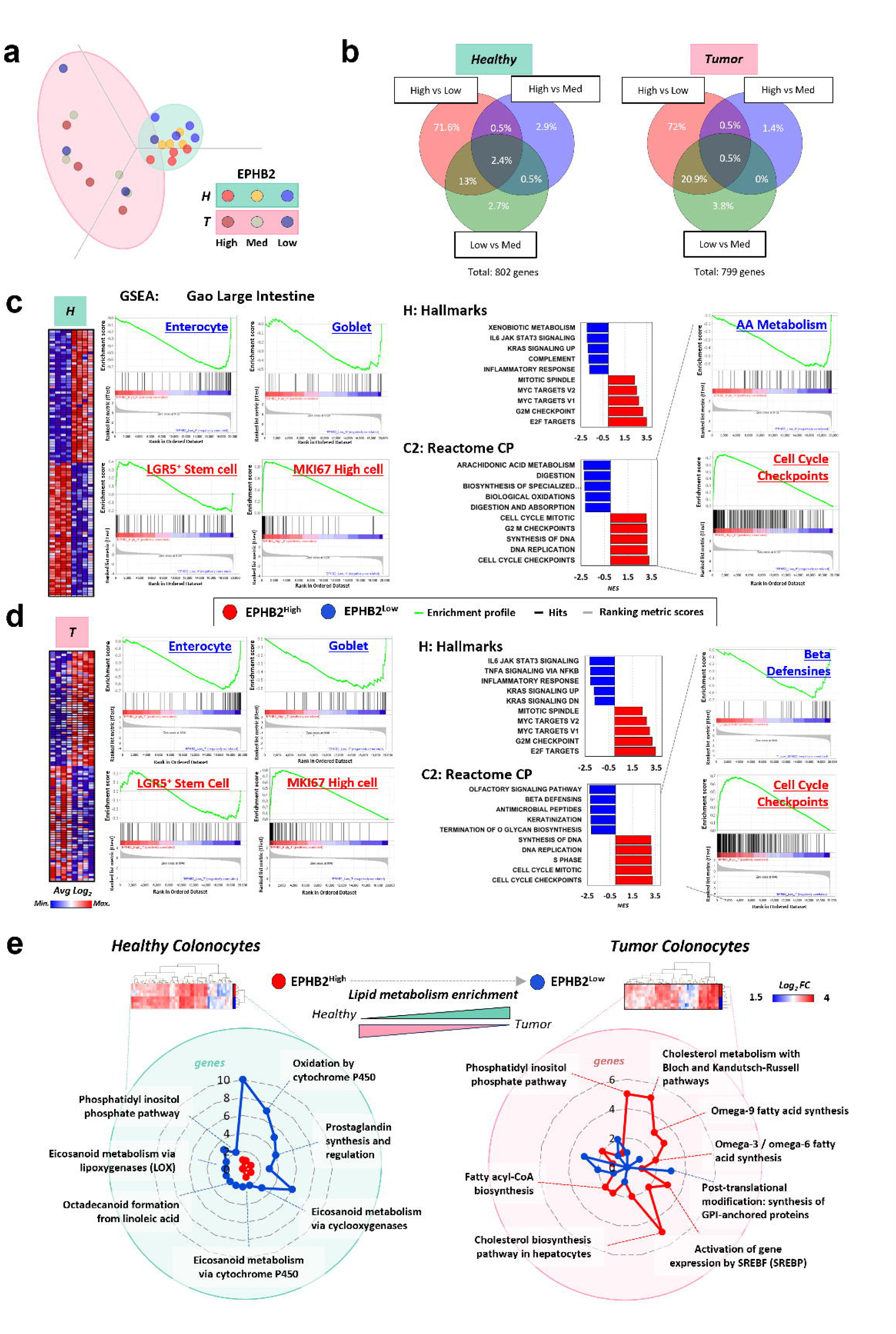
Transcriptomic profile of EPHB2 subpopulations showing gene expression of lipid-related enzymes is dependent on cell state in both healthy and tumor colonocytes. a) PCA analysis revealing a hierarchical distribution of the sample according to sample type (healthy or tumor) and EPHB2 subpopulations (Low, Med, and High). PCA was generated by mapping 73.3 % variance of CHP files (gene level information) with a maximum number of 3,000 data points. b) Venn diagrams of healthy and tumor EPHB2 subpopulations describing the percentage of genes differentially expressed according to FC ≥|2|, *P* ≤.05. c and d) GSEA of EPHB2^High^ vs. EPHB2^Low^ subpopulations in both (c) healthy and (d) tumor samples. The figure contains the heatmap of the top-50-ranked features for each subpopulation and enrichment for MSigDBs GAO large intestine: enterocyte, goblet, MKI67^high^, and LGR5^+^ signatures(31), H:Hallmark(32), and C2:Reactome curated datasets(33). (e) Unsupervised hierarchical clustering heatmap of differential expression analysis of lipid-related genes between EPHB2^High^ vs. EPHB2^Low^ subpopulations (FC ≥|2|, *P* ≤0.05, and presence of lipid-related genes Suppl. Table 5). Radial plot pathway enrichment analysis representing the number of genes overexpressed in each subpopulation (Wikipathways database(76), top-15 enriched pathways, 2.2 contingency in two sided Fisherʹs exact test, *P* <.05)

### Gene Co-expression analysis

R package for weighted correlation network analysis (WGCNA)(26) calculated the correlation patterns among genes expression samples EPHB2 High and EPHB2 Low subtypes, for both healthy and tumor samples, identifying highly correlated genes, and grouping into gene modules. First, a filtration step based on gene expression coefficient of variance of 0.05 was applied. Then, WGCNA parameters were set as unsigned bicor (biweight midcorrelation)(26) to calculate network adjacency prioritizing the absolute strength of the gene co-expression **(Figure 4b).** We selected soft-thresholding power to ensure the scale free topology, with a β-value of 12 (scale-free R^2^=0.90) for the healthy samples; and a β-value of 16 (scale-free R^2^=0.50-0.60) for the tumor **(Supplementary figure 13).** Minimum-module-size was set to 30 probes, and modules whose distance is less than 0.25 were merged. Intramodular hub-genes were identified based on combined gene-module membership and gene-trait significance scores. Detailed information of R code available at: https://github.com/idisba/lipid-RNA_Maimo-Barcelo.

**Figure 4.**
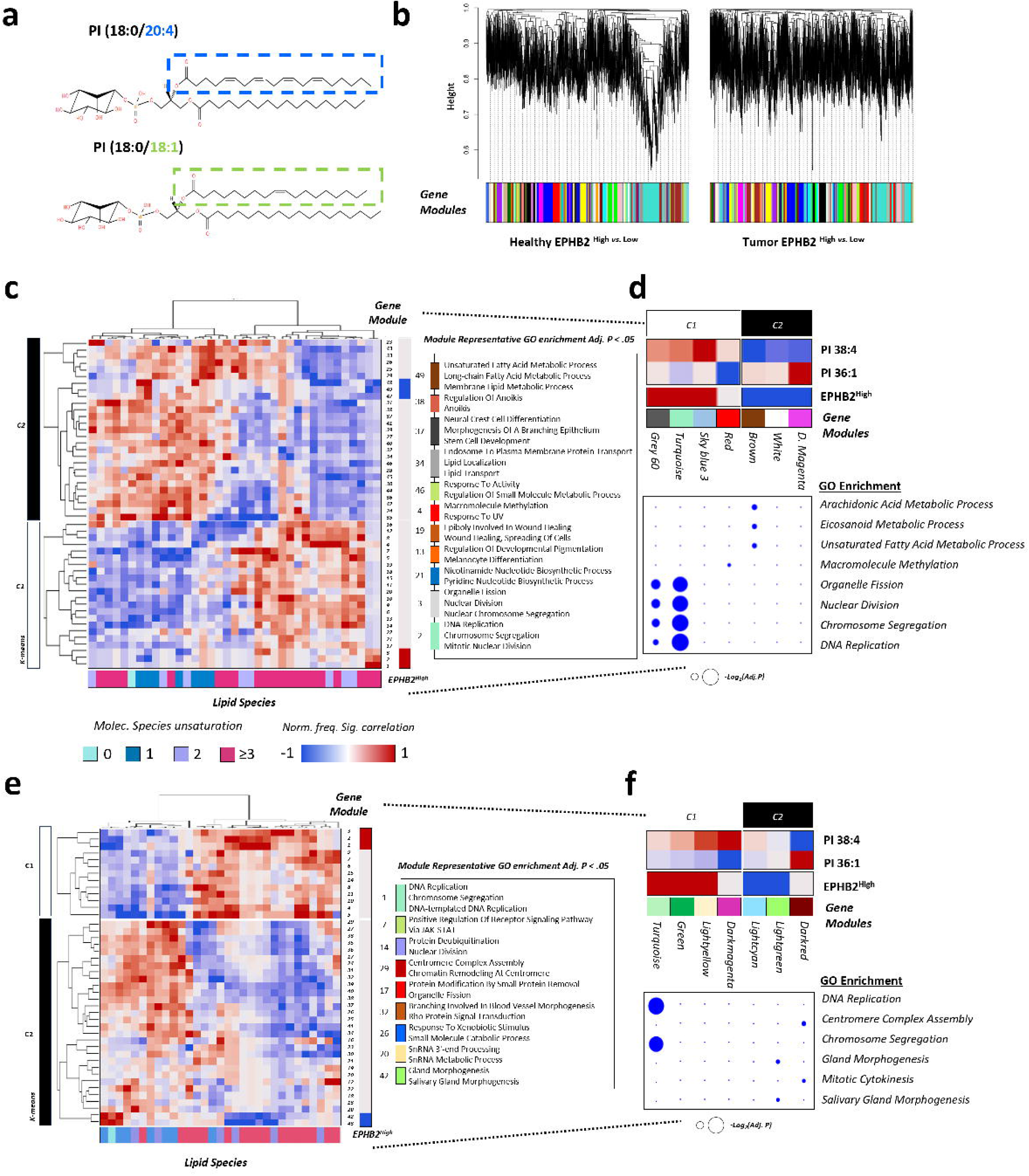
Lipidomic and transcriptomic data integration. a) Molecular structure of AA-PI 38:4, and oleic acid-PI 36:1 on sn-2 acyl chain position. b) R package for WGCNA was applied to transcriptomic data to identify modules of co-expressed genes based on their expression patterns. c-f) Lipid species were correlated with gene co-expression modules (CMs) associated with EPHB2^High^ vs. EPHB2^Low^ phenotype, to construct a novel integration matrix of lipid-RNA levels representative of colonocyte differentiation in healthy (c) and tumor (e) conditions (Pearson’s correlation coefficient, 576 permutations, *P* <0.05, lipid species color-coded according to molecular species unsaturation level 0 to >3). Representative GO terms overrepresented in CM are illustrated according to CM labels (Adj. *P*<0.05). CM with high correlation values for PI 38:4 and PI 36:1 profiles were selected along their GO enrichment dot plots (-log_2_(*Ajd*. *P*)) (d and f). K-means Cluster 1 = C1, Cluster 2 = C2 (c-f). List of GO enrichment of CMs is provided in Suppl. Table 8.

### Multiomics data integration

Obtained WGCNA gene modules were correlated using Pearson’s correlation coefficient with lipid species abundance of EPHB2 High and Low **(Supplementary table 7).** The correlation values were calculated for the maximum number of sample subtype permutations between lipids and gene-modules (n=4) (n_lipid_!×n_gene module_!). The number of significant correlations (P < 0.05) between each lipid and module was used to construct the final hierarchical heatmap correlation matrix (**Figure 4c-e**). Module gene ontology over representation analysis was calculated using R package clusterProfiler. Further network visualization and analysis of selected modules were performed with Cytoscape (v3.1) including STRING app to annotate Reactome pathway enrichment. Detailed information of R code available at: https://github.com/idisba/lipid-RNA_Maimo-Barcelo.

### Transcription factor enrichment analysis

The analysis was performed with iCisTarget(27,28) on the gene modules of interest from WGCNA. Modules were divided in two datasets according to kME sign. The official gene symbol was used as input data, and only datasets of ChIP-seq data (iCisTarget database) were interrogated. Parameters for analysis were set as default, briefly, region mapping was set from 10kb upstream of the transcription start site to 10kb downstream. Minimum fraction of overlap was set to 0.4 (40% of the input region or vice versa) and NES threshold set >3.0. The enrichment analysis within each database was made separately. Finally, region rank cut off or ROC threshold for AUC calculation was set to 0.005 to compare and rank all motifs and the threshold for visualization was set to 20,000 regions. Predicted transcription factors with NES ranked in top 3 were further network analyzed including target genes and calculating reactome pathway enrichment.

### Single cell RNA-Seq data

The GSE178341 dataset was analyzed in the single cell portal (singlecell.broadinstitute.org). Pearson’s correlation values were used to represent the similarity of each gene expression profile (row) with the *LGR5* profile and was used to order genes in a crypt-basal to apical fashion in **Figure 5h**. For **Figure 7**, we have visualized the gene expression colonocytes according to healthy (blue) and tumor (red) sample type condition, as well as for colonocyte subtype. Data was expressed in scaled mean expression (color scale) and % of expressing cells (dot size) in all cases.

**Figure 5.**
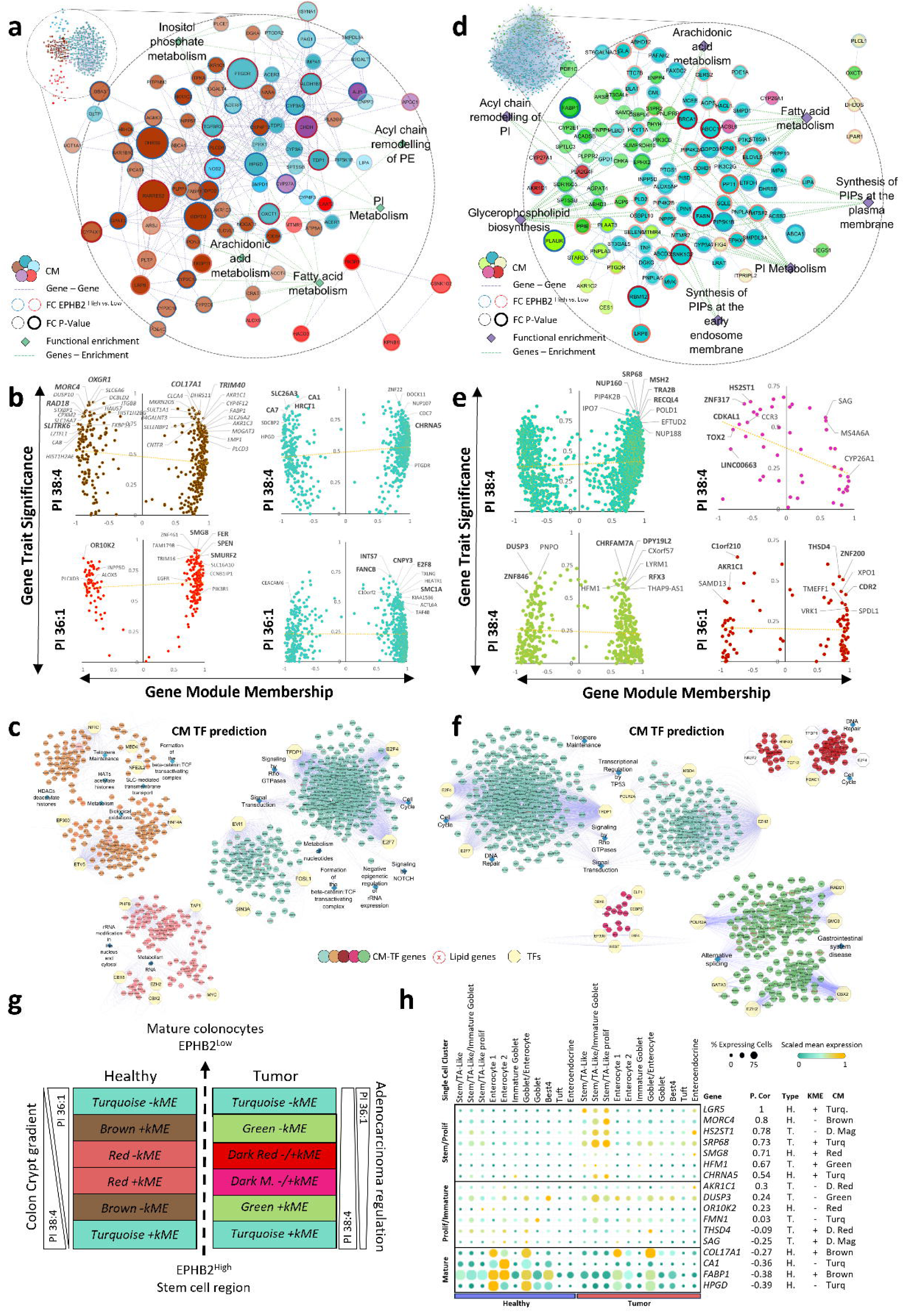
Gene expression network analysis of CM correlated with PI 38:4 and PI 36:1 phenotype in EPHB2 ^High *vs.* Low^ colonocyte differentiation healthy and tumor models. a) and d) Gene expression network of the top-5 CMs of PI 38:4 and 36:1 (small networks on the upper part) in healthy samples (a), and tumor samples (d). For both conditions, the zoomed networks contain only the lipid-related genes (92 genes in healthy, 130 genes in tumor), and their reactome functional enrichment analysis, calculated for each network and integrating the significant lipid-related processes. a) and d) Legend: Node color = co-expression module, node-size and border color = gene differential expression (blue, EPHB2^Low^; red, EPHB2^High^), green diamond = reactome functional enrichment process, edges = gene-interaction. b) and e) Representative plots of Hub gene analysis of CMs with the major contribution to PI 38:4 and PI 36:1 phenotype regulation. For each CM, the Top-5 genes with high combined score (|*kME_i_*|+ *GS_i_*) were highlighted as CM-PI hub-genes (bold-marked): In Healthy network, for PI 38:4, Brown CM (OXGR1, MORC4, RAD18, COL17A1, and TRIM40), and Turquoise CM (SLC26A3, CA1, CA7, HRCT1 and CHRNA5); for PI 36:1, Red CM (OR10K2, SMG8, FER, SPEN, and SMURF2), and Turquoise CM (INTS7, FANCB, CNPY3, E2F8, and SMC1A). In Tumor network, for PI 38:4, Turquoise CM (NUP160, SRP68, MSH2, TRA2B, and RECQL4), Dark Magenta CM (HS2ST1, ZNF317, CDKAL1, TOX2, and LINC00663), and Green CM (DUSP3, ZNF846, CHRFAM7A, DPY19L2, and RFX3); and for PI 36:1, Dark red CM (C1orf210, AKR1C1, THSD4, ZNF200, CDR2). c and f) Interaction network of top-3 TFs (octagonal nodes) with highest NES for each CM gene list according to KME sign. TF NES was calculated based on ChIP-seq data (iCisTarget), edges represent TF interaction with genes, for healthy samples (c) and tumor samples (f) analysis. g) Simplified model for gene CMs associated with the PI 38:4/PI 36:1 regulation during the colonocyte differentiation process. The model is based on EPHB2^High^ and EPHB2^Low^ isolated colonocyte gene expression profiles and organized according to KMEs of each CM. h) Dot-plot of single-cell gene expression analysis from human healthy and tumor colonocytes (GSE178341 dataset) (39). The gene expression of the top-combined-score-hub genes from healthy and tumor CMs (c and d), stem-cell marker *LGR5*, and mature colonocyte markers *FABP1* and *HPGD* were plotted to highlight the impact, and association of our calculated CM hub-genes in the different colonocyte subpopulations for both healthy and tumor colon cancer cells. Pearson’s correlation values represent the similarity of each gene expression profile (row) with the *LGR5* profile, and was used to order genes in a crypt-basal to apical fashion. Data were analyzed in the single cell portal (singlecell.broadinstitute.org).

### Organoid culture

Sample collection was approved by the Animal Ethics Research Committee IdISBa (2015, 17, AEXP). The protocol was adapted from Sato et al.,(29) using from crypt fragments from NMRI male mice. The mice colon was extracted, and the distal part (1 cm) was collected and flushed out with PBS+primocyn (0.2%) (InvivoGen, Spain). The tissue was minced into small pieces and washed twice in PBS with primocyn (0.2%) for 5 min. The tissue was resuspended in 5 ml of DMEM/F12 with 1% FBS (complete DMEM), and 500 U/ml collagenase and incubated at 37 °C for 30 minutes in a water bath shaking vigorously every 5 minutes. To finish dissociating the tissue and to stop the collagenase activity 5 ml of DMEM/F12 with 10% FBS was added and the tissue was mechanically dissociated with a 10ml-pipette. The top 5 ml of the resulting crypt suspension was transferred to a new tube, passing through a 100 μm cell strainer to remove large debris. The DMEM/F12 addition and filtration processes are repeated 3 times. Crypts were collected by centrifugation at 300×g for 5 minutes. Pellet was washed twice with 5 ml of complete DMEM, 60×g for 3 minutes, to reduce single-cell precipitation. Crypt fragments were mixed in 15 μl cold phenol red free Matrigel^TM^ matrix (Corning^TM^, Thermo Fisher Scientific, Spain) and plated in 96-well plate. The organoids were cultured with “WENR” culture media as described in Sato et al.(29): 50% Wnt and 10% RSPONDIN complemented media (kindly provided by L-Wnt3A cells and 293T-HA-RspoI-Fc cells from Hans Clevers lab), 40% DMEM F12 10% FBS, 1% primocin, and supplemented with B27, N2, 1mM N-acetylcysteine (NaC) (Thermo Fisher Scientific, Spain), 100 ng/ml noggin, and 50 ng/ml Egf (Peprotech, UK).

### PLA2-PTGS inhibition and prostaglandin treatments

The concentrations used were as follows: 5 µM for arachidonyl trifluoromethyl ketone (cytosolic PLA2 inhibitor), 20 µM for bromoenol lactone (Calcium-independent PLA2 inhibitor), 0.1 mM for valeroyl salicylate (PTGS1 inhibitor) and 4 µM for celecoxib (PTGS2 inhibition) (Cayman Chemical, USA). In the case of the PGD_2_ (Merck, Spain), PGF_2α_, and PGE_2_ (Bio-techne, Spain), all were used at 0.022 µM. The PTGER4 agonist L-902,688 was used at 1 µM while PTGER4 antagonist L-161.982 was used at 10 µM. All inhibitors were prepared following manufacturer instructions. Images of the organoids were obtained with an Axioscope Cell Observer microscope (Carl Zeiss, Germany) at 0 h, 24 h, and 48 h of treatment. The growth ratio was calculated by dividing the mean size of the organoids at 48h by the mean size of the same well at 0 h. The area quantification was done using Zen Blue Software (Carl Zeiss, Germany). The effect of the maximum concentration tolerated before organoid impairment for each inhibitor on organoid growth is shown in **Supplementary Figures 4 and 9**. No significant differences were observed on organoids growth and subtype marker expression under the synergic effect of the inhibitors in the absence of prostaglandins **(Supplementary figure 6)**.

The PLA2 and COX activities were assessed over isolated mice crypts after treatment, and was performed in the presence of the inhibitors at the working concentrations. In order to be able to compare all the samples, the protein content of each reaction was assessed as described in protein quantification methods. For PLA2 and PTGS inhibition assessment, the PLA_2_ activity was measured with the EnzCheck PLA_2_ Assay Kit of Invitrogen (Thermo Fisher Scientific, Spain) measuring the ratio between the emission at 515 and 575 nm. PTGS activity was measured with the COX Activity Assay Kit (Abcam, Spain), measuring the fluorescence emitted at 587 nm of a PTGS probe after its peroxidation **(Supplementary figure 5).**

### Organoid MTT assays

Colon organoids viability after the different treatments were assessed with Roche MTT assay kit (Sigma-Aldrich, Spain) following manufacture procedure. Synergy H1 microplate reader (BioTek, Germany) was used to measure colorimetric change. The signal was calculated by subtracting the reference absorbance to the 550 nm signal absorbance. The amount of absorbance is proportional to cell number.

### Digital Droplet PCR

The effects of prostaglandin signaling and its inhibition over colonocyte proliferation and differentiation was assessed quantifying the gene expression of different cell markers using digital droplet PCR (ddPCR). The RNA markers used were: *Krt20* for differentiated cells, *Kit* for the cKit^+^ cells, *Lgr5* for the stem cells and *Wdr43* for the TA cells. Total RNA was isolated from samples using Tripure reagent (Sigma-Aldrich) and following transcription to cDNA (up to 1µg RNA) with SensiFAST cDNA Synthesis Kit (Bioline, London, UK). Droplets were generated using QX200 droplet generator (Biorad) and the PCR reaction in a termocycler (C1000 Touch^TM^ thermal cycler, Biorad). The positive droplets were quantified using QX200 droplet reader (Biorad). The concentration values were used to assess differences between conditions. The PCR reactions were performed using QX200 ddPCR EvaGreen supermix, PCR Water Ultra Pure 18.2MΩ, DNase, RNase-Free (Bioline) and the following primers: Mouse *Lgr5*: Fw 5’-ACCCGCCAGTCTCCTACATC-3’, Rv 5’-GCATCTAGGCGCAGGGATTG-3’ (Tm = 57.5 °C); *cKit*: Fw 5’-TGGCTCTGGACCTGGATGAT-3’, Rv 5’- ATCTTTGTGATCCGCCCGTG3’ (Tm = 60.8 °C); *Krt20*: Fw 5’- TAGAGTTGCAGTCCCACCTC-3’, Rv 5’- TGTTCTTGGTTCTGGCGTTC-3’ (Tm = 60.8 °C); *Wdr43*: Fw 5’- AGTCCTCCTTACACAGGGCT-3’, Rv 5’- CGGTATCCTCAGCACAGTCC-3’ (Tm = 60.6 °C) (Isogen, Netherlands).

### Nuclei isolation from colonocytes, protein quantification and Western Blot

A pellet of isolated crypts was resuspended in 500 μl of PBS, sonicated (2 s, 10% amplitude), and further centrifuged at 100×g, 5 min, 4 °C. The supernatant was centrifuged at 1,500×g, 5 min, 4 °C, and the nuclei-enriched pellet was stored at −80 °C. Protein concentration was measured using a protein assay based on the Bradford dye-binding method, according to manufacturer’s instructions (Bio-Rad Laboratories, Madrid, Spain). Samples for western blot were mixed with electrophoresis loading buffer (120 mM Tris-HCl, pH 6.8, 4% SDS, 50% glycerol, 0.1% bromophenol blue, and 10% mM β-mercaptoethanol) and then boiled for 3 minutes. For the immunoblotting, equal protein content extracts were resolved in SDS-polyacrylamide gel and transferred to nitrocellulose membranes. Membranes were blocked with blocking solution (PBS containing 5% non-fat dry milk and 0.1% Tween) for 1h at RT. Then, the membranes were incubated overnight at 4 °C with the specific antibodies: rabbit anti-EP1 receptor (1:500, Bioss Antibodies, USA), rabbit anti-EP2 receptor (1:1500, Antibodies-online), rabbit anti-EP3 receptor (1:500, Bioss Antibodies), rabbit anti-EP4 receptor (1:500, Bioss Antibodies), rabbit anti-DP receptor (5μg/ml, Enzo Life Sciences, Spain), rabbit anti-FP receptor (5μg/ml, Enzo Life Sciences, Spain), rabbit anti Na^+^/K^+^-ATPase (1:10000, Abcam, UK) and β- actin (1:10000, LI-COR Biosciences, Spain). After the incubation, membranes were washed with PBS containing 0.1% Tween 20 (Sigma-Aldrich), and with 0.1% Tween 20 PBS containing 5% BSA (Sigma-Aldrich) for polyclonal antibodies. Then they were incubated with goat anti-rabbit IRDye 800CW (1:5000, LI-COR, Spain) or Alexa 680 donkey antimouse IgG (1:2500, Abcam) secondary antibodies at room temperature for 1 h. Membranes were visualized using an Odyssey CLx Imaging System (LI-COR). Quantity one software (Bio-Rad) was used to quantify the specific signals. Normalization was performed by the protein/β-actin content.

### Nuclear colocalization of prostaglandin receptors in colonocytes by Immunofluorescence

A biopsy fragment of 3 mm^3^ approximately from NMRI mouse colons was used for colocalization evaluations. To detect the presence of prostaglandin receptors in the nucleus, we immunolabeled each protein and analyze its colocalization with the nuclei labeled with DAPI using the Manders’ colocalization coefficient (MCC). Briefly, tissue sections were fixed with ice-cold methanol:acetone (1:1) for 10 minutes, and then were blocked with 5% BSA. Then, tissue sections were incubated with the following antibodies: rabbit anti-EP1 receptor (1:300, Bioss Antibodies), rabbit anti-EP2 (1:250, Antibodies-online), rabbit anti-EP3 receptor (1:400, Bioss Antibodies), rabbit anti-EP4 receptor (1:300, Bioss Antibodies), rabbit anti-DP receptor (1:500, Enzo Life Sciences), rabbit anti-FP receptor (1:500, Enzo Life Sciences) rabbit anti Na+/K+ ATPase (1:100 Abcam). Donkey anti-rabbit alexa fluor 555 (A31572, Life Technologies, 1:400) was used as secondary antibody and Fluoroshield™ with 4′,6-diamidino-2-phenylindole (DAPI) (Sigma-Aldrich, Spain) for nuclei. Images were obtained using Zeiss confocal microscope and the MCC were calculated using Zen software. Briefly, ROI were manually drawn around the nuclei (10 nuclei/crypt, 5 crypts section) in three consecutive sections. The MCC works as follows: for two probes (named R and G), two different MCC are derived, M1 is the fraction of R in compartments containing G, and the M2 would be the fraction of G in compartments containing R. These coefficients are calculated as follows: 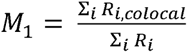 where *Ri* colocal = *Ri* if *Gi* > 0 and *Ri* colocal = 0 if *Gi* = 0, and 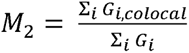 where *Gi* colocal = *Gi* if *Ri* > 0 and Gi colocal = 0 if *Ri* = 0. This method evaluates colocalization measuring co-occurrence regardless of the proportionality between the channels. To overcome the effect of the background over the MCC we used the point scatter of the colocalization graph to determine a good threshold for each channel (DAPI and the antibody). For each protein and image, a specific threshold was applied. Negative MCC values were assessed by analyzing tissue areas without nuclei but with membrane signal, and a negative area apart from the tissue. The threshold was established at the point where the protein channel was just above the negative colocalization areas and the increasing DAPI channel enough to have MCC values of 0 on the negative areas. To assess which MCC value indicate a true negative in the colon, we also calculate this coefficient for the Na^+^/K^+^-ATPase at the nucleus. Values equal to or below this exclusive cytoplasmic membrane protein MCC were considered negative.

## RESULTS

### Human colonocyte subpopulations membrane lipid profile recapitulates the gradient uncovered in colon tissue

Viable human healthy and tumor colonocytes were sorted based on EPHB2 expression levels, which correspond to the differentiation state of the epithelial cells(7), and labelled as EPHB2 High, Med (medium), Low, or Neg (negative) (**Supplementary Table 2**, n= 5-8, **Figure 1c**). The limited number of cells obtained from surgical biopsies posed a nontrivial challenge, which we overcame by developing a novel analytical strategy combining cell micro-deposition on glass slides with MALDI-MS (Figure 1d).(19) This approach provided higher sensitivity and facilitated comparison with tissue data.(11,23) Using this method, we achieved consistent and comprehensive lipid profiling from as few as 1.5-2·10^4^ colonocytes per subpopulation, a number 1–2 orders of magnitude lower than those commonly used in conventional lipidomic approaches.(30) Tumor subpopulations profiles differed from that of healthy subpopulations (**Figure 2a-b**). Thus, in tumor colonocytes, AA-containing PI species increased 1.5-fold, di-unsaturated fatty acids (DUFA)-containing PI species showed a 0.8-fold decrease, and MUFA-species levels increased 1.3-fold in PE while remaining similar in PI (**Figure2c-d**).

The most consistent lipid changes occurred between basal (EPHB2^High^) and apical (EPHB2^Low^) subpopulations (**Figure 2e-g**). The heatmap shows the hierarchical clustering of the top-ten differentially detected lipid species (**Figure 2f**), emphasizing the prominent role of PI species in colonocyte differentiation. In healthy colonocytes, AA-PI species (including PI 38:4 and PI 36:4)(11,23) content was higher in the crypt-base, EPHB2^High^ cells, than in the apical EPHB2^Low^ cells, while MUFA-PI species showed the opposite trend (**Figure 2h**). Consistent with our results in adenomatous polyps,(11) the difference between these species clearly diminished in tumor subpopulations (**Figure 2h**). Furthermore, AA-PI increased within tumor subpopulations, notably in the less differentiated tumor subpopulation, EPHB2^High^ cells (1.9-fold) (**Figure 2i**). On the other hand, the levels of PI 34:1, PI 36:1, and PI 36:2 reflected the gradual crypt distribution pattern seen in tissue MSI(11), following a slight upward and non-significant trend in healthy EPHB2^Low^ colonocytes compared to EPHB2^High^ (**Figure 2i**).

### Gene expression profile of colonocyte subpopulations is enriched in AA metabolism, prostaglandin synthesis and regulation, and PIPs metabolism pathways

PCA and hierarchical clustering analysis of the gene expression data clustered the samples according to the pathophysiological state of the colonocyte (**Figure 3a, Supplementary Figure 1a**). Subsequent analysis demonstrated that a direct comparison between EPHB2^High^ vs. EPHB2^Low^ subpopulations captured the vast majority (72%) of the changes in gene expression. Therefore, we streamlined the analysis by focusing on the EPHB2^High^ vs. EPHB2^Low^ comparison to highlight the changes associated with differentiation (**Figure 3b, Supplementary Figure 2**). Importantly, Gene Set Enrichment Analysis (GSEA) confirmed that the EPHB2-based sorting strategy yielded epithelial subpopulations that accurately reflected the cell types within the crypt hierarchy. We specifically probed for cell-type-specific gene signatures from the large intestine in the human Molecular Signatures Database (MSigDB).(31) EPHB2^Low^ cells showed high enrichment for mature colonocyte signatures (enterocytes and goblet cells), whereas EPHB2^High^ cells exhibited gene signatures linked to transit-amplifying (TA) proliferative colonocytes (MKI67^High^) and stem cell signatures (LGR5+, OLFM4^High^) (**Figure 3c**). Unlike their healthy counterparts, tumor EPHB2^High^ cells showed a lower LGR5+ enrichment, but higher OLFM4 enrichment signature, consistent with a disruption of the canonical differentiation sequence (**Figure 3d, Supplementary Figure 3a**).

Then, data sets were characterized using the human MSigDB collection hallmark gene sets (H: 55 gene sets)(32) and curated gene sets (C2:Reactome 1654 gene sets(33)) (**Figure 3c-d**). Both healthy and tumor EPHB2^High^ colonocytes were enriched in cell growth and proliferation processes. Conversely, EPHB2^Low^ were enriched in immune, inflammatory, and signaling processes. However, while healthy differentiated colonocytes were enriched in metabolic processes (xenobiotic, fatty acid, and bile acid metabolism), tumor-differentiated colonocytes were enriched in additional immune processes (interferon-gamma response, complement, and allograft rejection) (**Figure 3c-d; Supplementary Table 4**). The C2:Reactome datasets highlighted the significance of fatty acid metabolism in healthy differentiated cells, especially AA acid metabolism, which was absent in tumor-differentiated subpopulations (**Figure 3c-d, Supplementary Table 4**).

Finally, the results were filtered according to a comprehensive list of lipid-related genes (Supplementary Table 5). In healthy subpopulations, the EPHB2^High^ vs. EPHB2^Low^ comparison resulted in 54 differentially expressed lipid genes (FC >2, *p*-value <.05), 38 up- and 16 down-regulated in EPHB2^Low^. This lipid gene signature was sufficient to separate both subpopulations using hierarchical clustering (**Figure 3e**). Remarkably, the analysis revealed significant enrichment in differentiated colonocytes for eicosanoid metabolism and prostaglandin biosynthesis and regulation pathways, including genes such as *CYP4F12*, *AKR1C1*, *AKR1C2*, *CBR1*, and *HPGD* (**Figure 3e**). Conversely, in stem cell-like EPHB2^High^ colonocytes, only *PTGDR* was overexpressed. In tumor subpopulations, 47 lipid genes were differentially expressed: 21 up- and 26 down-regulated in EPHB2^Low^. In tumor EPHB2^High^ cells, phosphatidylinositol phosphate pathway genes were upregulated, including *PIP5K1A*, *PIP5K1C*, *PIKFYVE, PIK3R3,* and *PLCG1* (**Figure 3e, supplementary Table 6**). Further, genes involved in cholesterol-related pathways, and synthesis of n-9 and n-3/n-6 fatty acids (including *FASN*, *ACSL1*, and *ELOVL5)* were also upregulated. In contrast, tumor EPHB2^Low^ cells were less enriched in lipid metabolism pathways. It is worth mentioning that the differential expression analysis between tumor and healthy subpopulations (Supplementary Table 6) reinforced the alteration of AA metabolism regulation observed in healthy colonocytes, with significant downregulation of eicosanoid metabolism via cyclooxygenases, and prostaglandin synthesis and regulation pathways in tumor cells.

In summary, the phospholipid profiles of the subpopulations recapitulated the results and confirmed on-tissue AA-/MUFA-species gradients(11,12) and strengthened the link between PI composition and the maturation or pathological state of the colonocyte (**Figure 2**), while gene expression profiles highlighted the cellular states commonly found along the crypt: proliferative states at the base and differentiated states at the apical side (**Figure 3**), easing the integrative approach for multiomics data analysis.

### Omic data integration: uncovering the relationship between colonocyte co-expression gene modules and phospholipid species

Strikingly, the AA-/MUFA-PI shift involves the replacement of a single fatty acid, in many cases, AA by oleic acid (**Figure 4a**). Such precision points to regulatory pathways finely tuned to tailor composition at the species level. To gain insight into the transcriptomic program involved, we followed an integrative approach using the weighted gene co-expression network analysis (WGCNA),(26) which is a powerful method for uncovering unexpected patterns and connections that are easily overlooked using more targeted approaches.(34–38)

First, we identified the gene co-expression modules (CM) for EPHB2^High^ vs. EPHB2^Low^ comparison in healthy and tumor conditions, finding 49 and 43 modules, respectively (**Figure 4b**). Then, lipidomic data were integrated by calculating the correlation matrix between the CM eigengene and the normalized abundance of species of EPHB2^High^ vs. EPHB2^Low^ (**Supplementary Table 7, Figure 4c-e**). Correlation of the CM with the EPHB2^High^ phenotype surfaced a clear division: cluster 1 contained the modules positively correlating with EPHB2^High^, whereas cluster 2 included those with a negative correlation. Importantly, a similar clustering pattern was observed in tumor samples, indicating that the clustering faithfully conveyed healthy or malignant cell differentiation. Gene ontology (GO) enrichment analysis confirmed the presence of proliferative processes in the modules with a positive correlation in cluster 1 while cluster 2 was enriched in differentiation and metabolic processes (**Figure 4c and e**). Notably, cluster 1 was highly represented by species with ≥3 unsaturations, whereas species with fewer unsaturations were predominantly grouped within cluster 2 (**Figure 4c-e**). Thus, the clustering approach converged the unsaturation profiles and transcriptional landscapes across differentiation stages.

Moreover, during healthy differentiation, correlation values highlighted the clear inverse relationship between AA- and MUFA-containing PI at the module level (**Figures 1b and 4c**). Thus, the proliferative modules of cluster 1 were positively correlated with the PI 38:4 profile, while cluster 2 modules were correlated with PI 36:1 (**Figure 4d**). Among these modules, the brown one, with the greater negative correlation for PI 38:4, had significant GO enrichment for AA, eicosanoid, and unsaturated fatty acid metabolic processes. Similarly, tumor colonocytes also showed the opposite module correlation profile for AA and MUFA-PI species with a strong presence of cluster 1 modules (**Figure 4f**). Nonetheless, the modules showing a high correlation for PI 38:4 and 36:1 (dark red and dark magenta, respectively), were not enriched for AA metabolism (**Figure 4f**).

### Network analysis and transcription factor (TF) prediction of integrated data

To better understand the transcriptional control of PI gradients, the gene composition of the CM involved in PI 38:4 and 36:1 regulation was analyzed (**Figure 6d and f**). Thus, gene interactions were visualized as a network to gain insight into the polarized module distribution during differentiation for healthy and tumor samples. In healthy colonocytes, the network was composed of the top-five PI 38:4 and 36:1 CM (turquoise, skyblue3, red, brown, and dark magenta (**Figures 4d and 5a**). The modules contained at least 92 genes related to lipid metabolism, for which the functional enrichment analysis retrieved processes specific to membrane lipids, such as PI and inositol phosphate metabolism, as well as AA and fatty acid metabolism, all tightly related to PI 38:4 and 36:1 regulation (**Figure 5a**). We selected the brown, red, and turquoise modules for further analysis due to their higher content of regulatory lipid genes.

**Figure 6.**
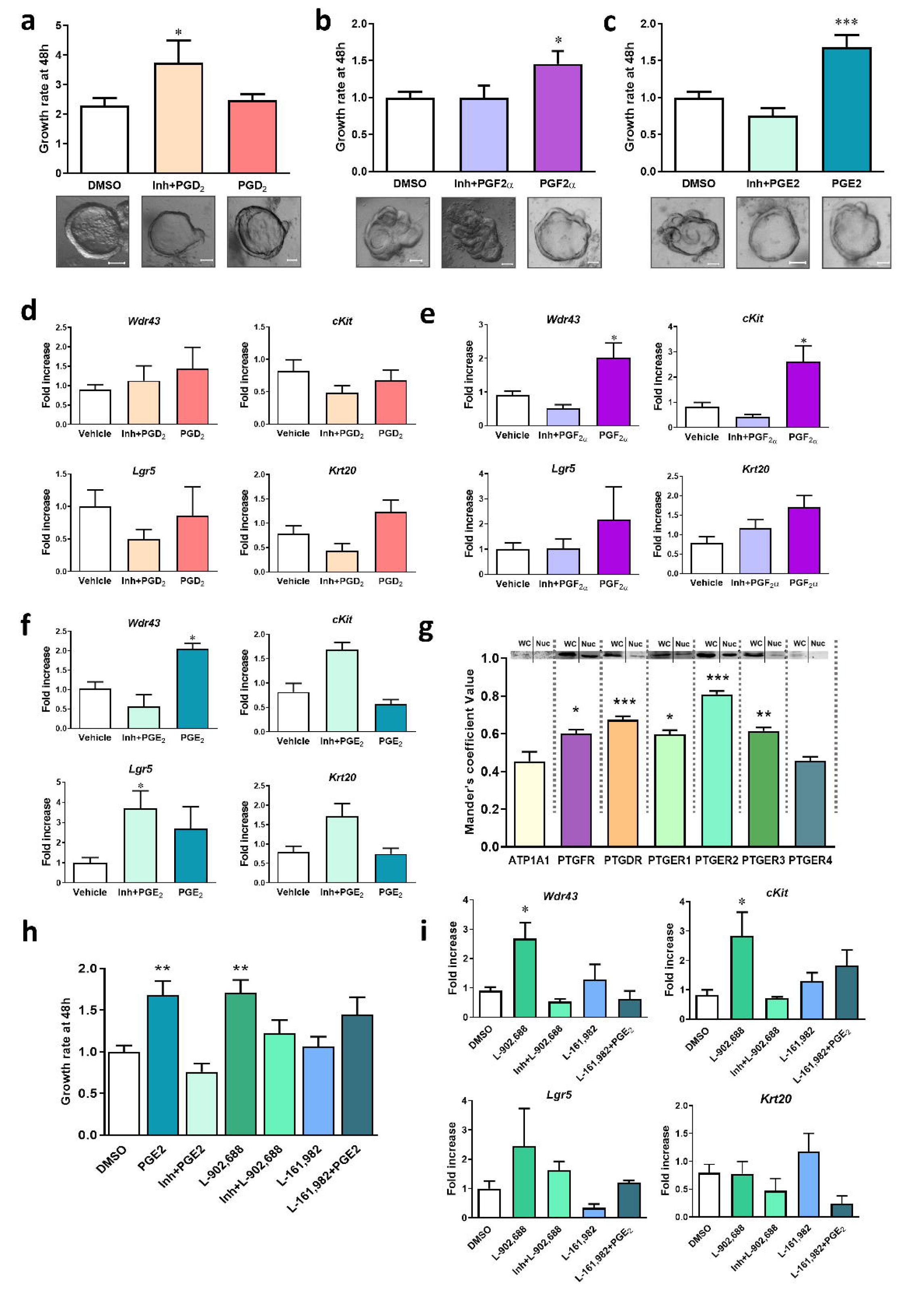
Effects of major prostaglandins on murine colon organoid proliferation and differentiation. a-c) Bar plots showing the effects on organoid growth ratio, and representative images of organoids for each treatment. Scale bar = 50 μm. Organoids were treated with PLA2, PTGS1 inhibitors, and with PGD_2_ (a), PGF_2α_ (b), or PGE_2_ (c); d-f) Changes in gene expression: Wdr43 (TA cells), cKit (Paneth-like cells), Lgr5 (Stem cells), and Krt20 (enterocytes) upon treatment with PGD_2_ (d), PGF_2α_ (e), or PGE_2_ (f) and PLA2 and PTGS inhibitors. g) Mander’s coefficient value of prostaglandin receptors and representative Western Blot in crypts “C” or nucleus enriched fraction “N” (n=3). h) Effects of PTGER4 on organoid proliferation and differentiation. Organoids were treated with vehicle (DMSO), PTGER4 agonist L-902,688 (10 µM) alone and in combination with PLA2 + PTGS inhibitors, PTGER4 antagonist L-161,982 (10 µM), and PGE2 + L-161,982 (n=8). i) Impact of treatment with EP4 agonist and antagonist on colonocyte subtype differentiation. In all plots, values represent the mean ± SEM referred to control treatments (DMSO) (n=3-8). Statistical differences were established using one-way ANOVA followed by Bonferroni post-test, *\*p <*.05, \*\**p* < .001, \*\*\**p* <.0001.

**Figure 7.**
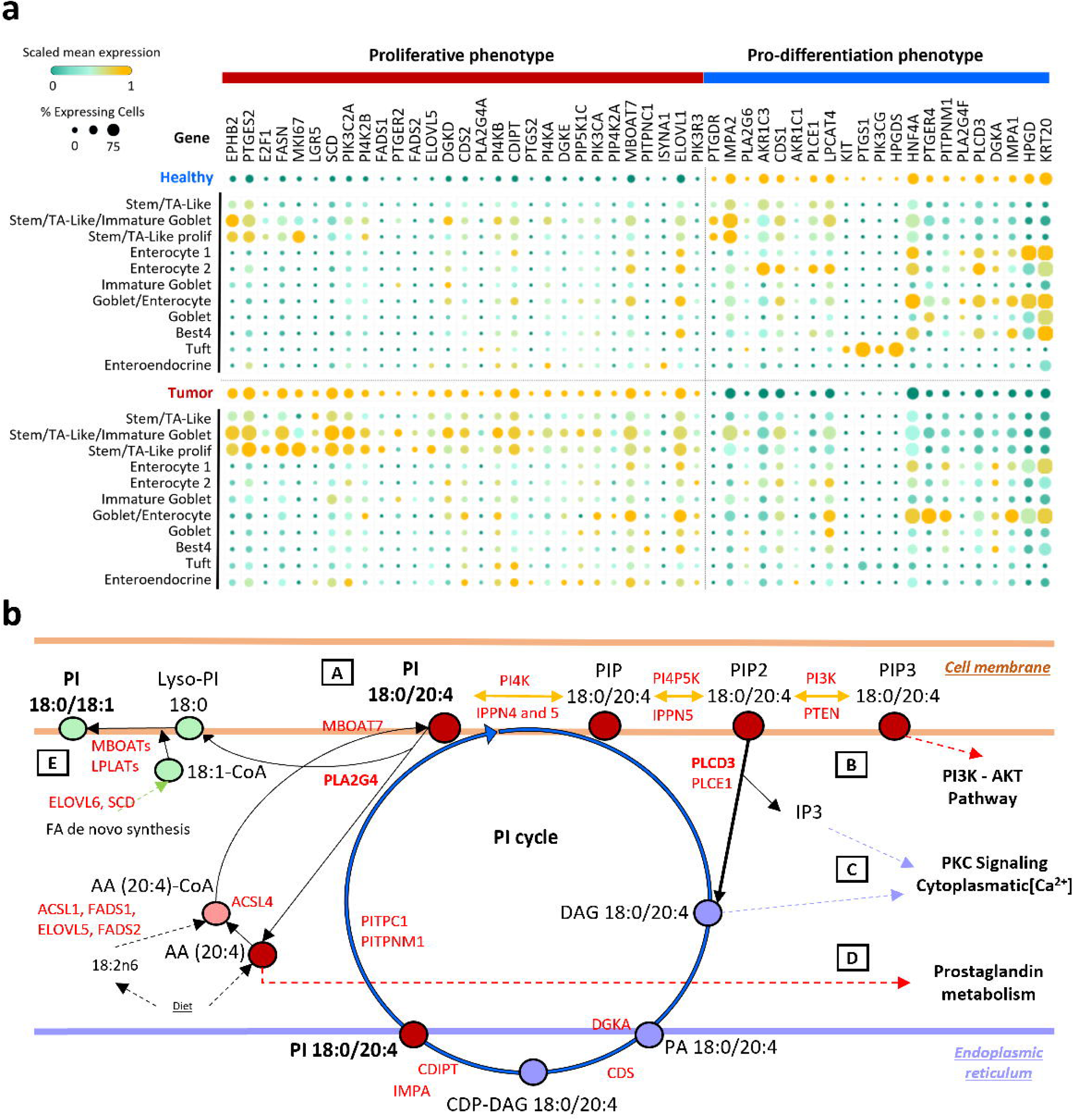
PI 38:4 metabolism in colonocyte differentiation and its deregulation in tumor colonocytes. a) Dot plot of Single-cell RNA-Seq data (GSE178341 dataset)(39) from human healthy and tumor colonocytes. The data show the gene expression of PI metabolism associated genes, including: PIP biosynthesis and PI cycle, supply and biosynthesis of AA, and synthesis and esterification of 18:1 n-9 in PI. Gene expression is illustrated into two categories: sample type healthy and tumor rows, and colonocyte subtypes (from stem cell and TA-like to the different mature and specialized subtypes: enterocyte, goblet, best4, tuft, and enteroendocrine). In both categories, tumor colonocyte shows the proliferative phenotype of PI metabolism with upregulation of the metabolic genes of AA and PIPs biosynthesis in stem and TA subtypes, while healthy colonocytes show pro-differentiation phenotype with upregulation of inositol phosphate and prostaglandin metabolism. Colonocyte markers EPHB2, LGR5, KIT, and KRT20 as well as TFs E2F1 (proliferative CM) and HNF4A (healthy metabolic CM) are included as reference genes to facilitate the link the previous figures on the manuscript. Data were analyzed in the single cell portal (singlecell.broadinstitute.org). Upregulation was calculated according to reactome database (P<0.05). b) Pathway illustration of PI 38:4 metabolism in colonocytes physiopathology: steps “A” to “E” (black boxes) represent the metabolic remodeling of PI 38:4 shift to PI 36:1, including: PIPs synthesis (steps A to B) (genes: PI4K, IPPN4 and 5, PI4KP5K, IPPN5, PI3K, PTEN); PI cycle: DAG and IP3 generation (B and C) (genes: PLCD3, PLCE1, DGKA, CDS, CDIPT, IMPA, PITPC1, PITPNM1); AA synthesis, esterification and hydrolysis, and prostaglandin synthesis (step D) (genes: ACSL1, ELOVL5, FADS1 and 2, ACSL4, PLA2G4, ACSL4, MBOAT7, prostaglandin synthesis and regulation programs); and Oleic acid esterification into Lyso-PI 18:0 (step E) (ELOVL6, SCD, MBOATs, LPLATs). Red names, lipid enzymes; Oleic acid metabolism, green dots; AA in PI and PIPs metabolism, red dots; AA in DAG, PA and CDP-DAG in violet dots; cell membrane, orange stripes; endoplasmic reticulum in violet stripe. PI 36:1 and PI 38:4 are marked in bold.

Further, genes can also be distributed according to gene-trait-significance of PI 38:4 or PI 36:1 and gene-module-membership (kME)(26), which measures gene intramodular connectivity **(Figure 5b, Supplementary Table 8**). Thereby, in the brown module and PI 38:4 phenotype, *COL17A1* and *TRIM40* were identified as driver genes with positive membership. Their expression was upregulated in the crypt apical side (EPHB2^Low^ cells), and the genes found in positive kME were part of the aforementioned AA and PI metabolic processes (**Figure 5a-b**). On the negative side of the plot, *OXGR1, MORC4, RAD18,* and *SLITRK6* were identified as driver genes, which were found more expressed at the basal part of the crypt and functionally involved in processes of HDAC deacetylate histones, HAT acetylate histones, and DNA methylation (**Figure 5c**).

Finally, we looked for TF binding motifs significantly overrepresented in co-expression module signatures (ChIP-Seq TF binding sites, iCisTarget(27)) (**Figure 5c-f, Supplementary Table 9**). The results for the brown module identified ETV5, HNF4A, and EP300 as regulators of -kME genes, and MBD4, NFIC, and NFE2L2 for +kME genes (**Figure 5c**). The co-expressed module genes called by these top enriched TFs were plotted as a network, and the reactome functional enrichment annotations highlighted the similarity between the overall CM and the TF-filtered functional enrichment of CM-genes, emphasizing their functional regulation (**Figure 5c, Supplementary Table 9**).

The turquoise module expression profile displayed higher gene trait significance for PI 38:4 than PI 36:1, with hub-genes *SCL26A3, CA1, CA7*, and *HRCT1* for -kME (apical-crypt-side), while CHRNA5 was identified for +kME (**Figure 5b**). The latter contained most of the genes related to strong enrichment in cell division processes; thus, TF analysis identified E2F4, TFDP1, and E2F7 in +kME. The results for -kME TFs were *SIN3A*, *FOSL1*, and *EVI1*, whose associated genes were more related to apical crypt signal transduction, including the 15-hydroxyprostaglandin dehydrogenase, *HPGD,* and inositol phosphatase, *IMPA1* (**Figure 5c**). In the red module, a negative regulator of PI 36:1 crypt phenotype, the driver genes were *SMG8, FER, SPEN,* and *SMURF2* for +kME, and *OR10K2* for -kME (**Figure 5b**). TF analysis identified PHF8, TAF1, and MYC in +kME, regulating the expression of genes involved in RNA metabolism and rRNA modification, especially genes related to MYC; whereas in -kME enrichment was for EZH2, CBX2 and CBX8 (**Figure 5c**). Genes from +kME enriched in PHF8 and CBX8 were related to the regulation of genes associated with the regulation of phosphoinositide metabolism (*MTMR1* and *PIK3R1)*, and AA metabolism (*ALOX5),* respectively.In tumor colonocytes, five CM were used to construct the PI network (Turquoise, green, dark magenta, dark red, and light-yellow, **Figures 4f and 5d**), containing more than 130 lipid-related genes, with the turquoise and green modules containing the largest number. The gene network was enriched in processes such as glycerophospholipid biosynthesis, PI, PIP, and AA metabolism, especially in turquoise and green modules (**Figure 5d**). Remarkably, the dark magenta and dark red modules, which showed the highest correlation values for PI 38:4 and 36:1 (**Figure 4f**), contained two (*ACSL6* and *CYP26A1*) and three (*PLA2G4F*, CYP27A1, and *AKR1C1*) lipid genes, respectively (**Figure 5d**). Thus, we selected these last four modules to study their gene composition. Gene trait significance of the turquoise module was higher for PI 38:4 than PI 36:1(**Figure 5e**). We identified *SRP68, NUP160, MSH2, TRA2B,* and *RECQL4* as driver genes, all with positive module membership (+kME) and higher expression in EPHB2^high^ tumor cells. Remarkably, E2F4, TFDP1, and E2F7 were highlighted in TF analysis, as in healthy samples; while -kME genes predicted EZH2, MBD4, and POLR2A, associated mostly with signal transduction processes (**Figure 5e-f**). In the green module, *DPY19L2, CHRFAM7A,* and *RFX3* were identified as driver genes in +kME. Top predicted TFs were CBX2, EZH2, and GATA3. Finally, for -kME genes, *DUSP3* and *ZNF846* were the driver genes and POLR2A, RAD21, and SMC3 were the predicted TFs. Both kME were related to alternative splicing and gastrointestinal system disease processes (**Figure 5e-f**).

The combined score of the dark red module – with the best correlation with PI 36:1 in tumor samples – was similar for both kMEs. *THSD4, ZNF200,* and *CDR2* were identified as +kME hub-genes, while *C1orf210* and *AKR1C1* were -kME driver genes. The predicted +kME TFs were TFDP1, E2F4, and HDAC1, mostly associated with cell cycle and DNA repair; whereas the - kME TFs were HNF4G, TCF12, and NR2F2 (**Figure 5e-f**). Finally, all the top-score driver genes were identified in -kME: HS2ST1, ZNF317, CDKAL1, TOX2, and LINC00663 in dark magenta. The predicted TFs were CBX8, CEBPB, and ELF1. Regarding +kME genes, the predicted TFs were REST, EP300, and PCR1X. Such a small network did not retrieve significant results for functional analysis (**Figure 5e-f**).

Overall, the network analysis and TF prediction of integrated data strongly highlighted the association between specific gene modules, lipid species (PI 38:4, PI 36:1), TFs, and biological processes in both healthy and tumor samples. The changes observed in both transcriptomic and lipidomic profiles provided new insights into the regulatory mechanisms underlying colonocyte phenotype and their dysregulation in tumorigenesis, involving alterations in genes involved in PI regulation and prostaglandin metabolism. To better visualize the gradual regulation of PI 38:4 and PI 36:1 in our integrated data, the results were organized on a gradient from high to low EPHB2 expression, based on kME values (**Figure 5g**). This gene expression model highlights the spatial contribution of each CM, illustrating the shift in PI 38:4 and PI 36:1 during the transition from stem/progenitor cells to differentiated colonocytes. This model was further validated in a single-cell RNA-seq dataset of human healthy and tumor colonocytes (GSE178341)(39) using the hub-genes of each CM (**Figure 5h**), reinforcing the proposed regulation networks based on integration data.

### Prostaglandin metabolism in healthy colon organoid growth and differentiation

#### Effect of prostaglandin stimulation on colon organoid growth and differentiation

Transcriptomic analysis strongly reinforced the implication of prostaglandin synthesis and regulation during healthy differentiation, which aligns with the changes in AA-containing phospholipids in EPHB2 subpopulations and tissue sections.(11,12) To study the potential link between AA and eicosanoid metabolism and colonocyte differentiation, we administered PGD_2_, PGF_2α_, and PGE_2_ to developing colon organoids. In addition, we inhibit the activity of phospholipases with high AA affinity as well as PTGS1 and PTGS2, the key enzymes necessary for the production of prostaglandins from AA. We assessed the effects of prostaglandins and enzymatic inhibition on organoid growth and cell composition, by quantifying the expression levels of colonocyte subtype-specific markers: *Lgr5* for stem cells, *cKit* for Paneth-like cell, *Wdr43* for transit-amplifying TA cells, and *Krt20* for differentiated colonocytes,(40,41) using digital PCR. Additionally, organoid size was assessed as a proxy for proliferation alterations and cell survival (**Supplementary Figures 4-6**). Treatment effects were prostaglandin-dependent. Thus, PGD_2_ showed no impact on organoid size. However, PGD_2_ together with PLA2 and PTGS inhibition, resulted in a 1.6-fold increase in organoid size, while expression levels of the selected markers were not affected (**Figure 6a**). Conversely, PGF_2α_ increased organoid size 1.7-fold compared to controls, coinciding with a marked upregulation of Paneth-like and transit-amplifying cells (2.6- and 2.0-fold, respectively). PLA2-PTGS inhibition reverted the specific PGF_2α_ effect on proliferation (**Figure 6b**). Likewise, PGE_2_ increased organoid size 1.7-fold, which correlated with TA cell overexpression (2.0-fold). However, PLA2-PTGS inhibition prevented PGE_2_-induced changes on organoid size and *Wdr43* overexpression, leading to remarkable *Lgr5* overexpression (3.7-fold) (**Figure 6c and f**).

#### Role of prostaglandin receptors in colon organoid growth and differentiation

The effects of the different prostaglandins on colon organoids and their modulation under biosynthetic inhibition raised the question of how, the localized production of prostaglandins contributes to colonocyte differentiation. Thus, we investigated the nuclear and cytoplasmatic localization of the prostaglandin receptors PTGFR, PTGDR, and PTGER1-4 in colonocytes. Except for PTGER4, all receptors showed nuclei-colocalization values, which pointed to pathways that are more complex than initially expected (**Figure 6g and Supplementary Figures 7-9**). To assess whether changes in *Lgr5* expression occurred via PTGER4, organoids were incubated with the PTGER4 agonist (L-902,688) and antagonist (L-161,982). Similar to the PGE_2_ treatment, L-902,688 increased organoid size 1.7-fold, while PLA2-PTGS inhibition reversed this effect (**Figure 6c and h**). L-161,982 did not change organoid growth rate but blocked PGE_2_ effect on organoid growth (**Figure 6h**). Conversely, L-902,688 not only increased organoid size at a similar rate as PGE_2_ but also *Wdr43* and *cKit* expression (**Figure 6i**).

Altogether, exogenous PGF_2α_ and PGE_2_ showed a major impact on the basal region of the crypt, favoring colonocyte proliferation through stimulation of TA cells and inducing increased organoid size. Moreover, exogenous PGF_2α_ was also involved in *cKit* overexpression, suggesting a wider scope of impact. Importantly, the effect of exogenous supplementation relied on the enzymatic availability of PLA2-PTGS. PGE_2_ favored the overexpression LGR5-stem cell marker under PLA2-PTGS pharmacological inhibition without increasing organoid size (**Figure 6c and f**). Likewise, the effect of PGD_2_ on organoid growth/size also depended on PLA2-PTGS inactivity (**Figure 6a and d**), resulting in a pronounced increase in organoid size. These results strongly reinforce that PTGER4 activation contributes to the PGE_2_-dependent increase in colonocyte proliferation (**Figure 6f**).

## DISCUSSION

The rapid advances in MS-based analytical techniques applied to lipids exposed the stunning lipid diversity occurring in biological samples. The irruption of spatially resolved lipidomic techniques added complexity, demonstrating that the lipidome is cell type-specific and also highly sensitive to changes in the pathophysiological state.(18) Unfortunately, knowledge of how cells orchestrate this lipid repertoire continues to lag behind because of lipid metabolism complexity, which hinders the study of the myriad of lipid species coexisting in cell membranes.(17) In this context, colon epithelium MSI images show tens of phospholipid species changing in an orderly and coordinated manner along the crypt, from adult stem cells up to fully differentiated colonocytes,(11,12,23,42) strongly indicating that membrane lipidome regulation must be a crucial checkpoint in deciding stem cell-fate. Herein, we aimed to shed some light on the underlying regulatory mechanisms associated with the gradual change in lipid species and cell differentiation.

The integrative data-driven modeling of MSI and gene expression results enabled the establishment of a clear correlation between gene CM and lipid species involved in colonocyte stemness and maturation state. Particularly, this approach allowed the generation of gene networks modelling the mechanism underlying strict PI 38:4 distribution and the involvement of canonical pathways acting during colonocyte differentiation and oncogenic transformation (**Figures 4 and 5**).

In scenario of the differentiation, we identified the CM enriched in PI and AA metabolism (brown) and strongly correlating with PI 38:4 content, and the proliferative module (turquoise) linked to EPHB2^High^ cells. The brown module included phospholipases (PLA2G4F, PLCD3, PLCE1), inositol phosphatase (INPP5F), DGKA, and prostaglandin enzymes (AKR1C1, AKR1C2, AKR1C3) which, in combination, could act as a key program for AA mobilization and led to lower PI 38:4 levels. In agreement with literature, PLCD3 is a highly relevant candidate for PI profile regulation at the enzymatic level. Its overexpression in differentiated colonocytes (**Figure 5**) agrees with its function in microvilli formation of enterocytes in mice.(43) Furthermore, PLCD3 activity impacts cell proliferation capacity by generating DAG and IP3 from PIP_2_, leading to calcium increase and PKC activation,(44) and acting as a negative regulator of PI3K-AKT activation via PIP_2_ hydrolysis.(44) PLCD3 activation is driven by exogenous signals, such as the non-canonical Wnt pathway, via stromal-secreted WNT5A(45), and dietary vitamin D3, contributing to the calcium gradient along the crypts.(46–48) Besides, downstream activation of the PKC-MAPK pathway mediates enzymatic PLA2G4F activation,(49) which is key for AA-hydrolysis and eicosanoid synthesis. Finally, AKR1C isoforms 1, 2, and 3 participate in the biosynthesis of PGF_2α_ and regulation of PGD_2_ that, according to our results in organoids, play different roles in proliferation, with PGF_2α_ strongly linked to this process (**Figure 6**). Interestingly, the brown module also contained hub-genes implicated in important processes in colonocyte pathophysiology; such as COL17A1, involved in human colon cancer stem cell (LGR5^+^ p27^+^) dormancy(50); OXGR1, a GPCR that increases intracellular calcium upon α-ketoglutarate binding(51) (**Figure 5**); and EMP1, associated with residual EMP1+ tumoral cell population, which plays a role in metastatic recurrence in colon cancer.(52) Therefore, this module also seems to have an impact on colonocyte stemness. Moreover, the brown module showed a significant motif overrepresentation for TF HNF4A (**Figure 5**), a key driver of the colonocyte subtype differentiation and cell apoptosis,(53) strengthening the link between the regulation of PI remodeling and colonocyte fate.

Conversely, the healthy proliferative module bears three genes clearly linked to PI remodeling: ISYNA1, a rate-limiting synthetic enzyme for inositol-containing compounds,(54) critical in the generation of the PI 38:4 pool; PTGDR, a key receptor in PGD_2_ signaling,(55) linking prostaglandin metabolism with stem-cell compartment (**Figure 6**); and *HPGD*, a prostaglandin catalyzing enzyme, highly expressed in differentiated colonocytes and downregulated in tumor cells.(13,56) Overall, our results in colon organoids revealed the diverse impact of PGE_2_ signaling on colonocyte-subtype proliferation. Early studies showed that PTGER2 promotes β-catenin nuclei translocation during the initial stages of colonocyte proliferation,(57,58) while PTGER4 acts during intestinal wound healing and in mucosal protection.(59) Consistently, PTGER4 activation by agonists increased TA and Paneth-like cell markers in colon organoids (**Figure 6**). PGF_2a_ and PGD_2_ synthesis by mature colonocytes could contribute to the PGE_2_ negative regulation suggesting an autocrine signaling mechanism along the crypt, regulating cell proliferation. Within mature enterocytes, tuft cells shows high expression of *PTGS1* and *HPGDS,* key in PGD_2_ synthesis, while tuft cell marker *DCLK1* expression decreases in colon cancer (**Figure 7, Supplementary figure 11**),(39) consistent with PGD_2_-mediated antitumor role of tuft cells in pancreatic and gastric cancer.(60–62) Therefore, our results point to a prostaglandin metabolism dependent on cell-state and colonocyte subtype.

Our data also show differential expression of the prostaglandin receptors in the nuclei of colon epithelial cells (**Figure 6g**). In this sense, PTGS1 and 2 can be also localized at the nuclear envelope reinforcing the idea of a nuclear prostaglandin signaling(63). The role of these receptors at nuclear level is poorly understood, however, the presence of calcium mobilization mechanisms and phosphoinositide signaling in the nucleus could suggest a similar mechanism of action than at cytoplasmic membrane levels(64,65). For instance, nuclear PTGER3 stimulation induces calcium mobilization that, in turn, modulates eNOS expression in a PI3 kinase/Akt and Erk-MAP kinase pathways-dependent(66). Furthermore, PTGER2 nuclear signaling have been described associated to the differentiation state of small intestine epithelial cells. Thus, PTGER2 can be found in the undifferentiated epithelial cells while differentiated cells present only cytoplasmic expression. Nuclear PTGER2 expression by these cells would be associated to a genotoxic protective role(66). Thus, the functional PTGS-PTG coupling to the receptor at nuclear level would influence stem cell/proliferative pathways. Indeed, depolarization of the nuclear membrane by Ca^2+^ is a key step in nuclear translocation of β-catenin, one essential signaling feature of intestinal stem cells(66).

In tumor cells, the WGCNA analysis linked the altered PI profile to the aberrant proliferation program. The tumor proliferative module (turquoise) was the major transcriptomic program in terms of gene number, whereas the metabolic-like-module (i.e., healthy brown-like) was lost (**Figures 4 and 6**). The expression of enzymes mediating PI 38:4 metabolism during tumor colonocyte differentiation reflected a metabolic reprogramming affecting PI remodeling and PIPs synthesis. This was further reinforced at the level of hub-genes, showing *PIP4K2B* (phosphoinositide synthesis) among the top-ten genes for PI 38:4 phenotype. It is worth stressing that, although the participation of PI3K-AKT signaling in tumor progression is well established,(67) the specific regulation of PI species contributing to the phosphoinositide pool remains elusive. Recent in vitro studies show PI 38:4 favors PI channeling of into PIPs and facilitates its recycling to PI (through DAG–>PA–>CDP->DAG).(68–70) Herein, we show that PI 38:4 increased levels in tumor cells (**Figure 2**) was coupled to enrichment in PIP synthesis genes, and a lower maturation state (**Figures 4-6**). A plausible explanation is that tumor cells could maintain high PI 38:4 levels by sustaining high PI recycling activity and avoiding the upregulation of differentiation driving enzymes, such as PLCD3 and PLA2G4F (**Figure 7**). Thereby, we believe the maintenance of high PI 38:4 levels fuel the PI3K/AKT pathway, promoting and sustaining cell proliferation and a non-differentiated cell state. Although experimental validation falls beyond the scope of the present study, the proposed PI 38:4–PI3K/AKT connection is firmly grounded in established biology. PI serves as the universal precursor for all signaling phosphoinositides, including PI(4,5)P₂ and PI(3,4,5)P₃, the latter being indispensable for AKT membrane recruitment and activation via its pleckstrin homology domain.(66) Sustained availability of PI species enriched in the canonical stearoyl–arachidonoyl (18:0/20:4) acyl chains has been shown to favor the generation and stabilization of signaling-competent phosphoinositides. Moreover, AKT activity is strictly restricted to membranes containing PI(3,4,5)P₃ or PI(3,4)P₂, where membrane engagement not only enables phosphorylation but also protects AKT from dephosphorylation, thereby prolonging signaling duration.(66) Importantly, lipid acyl chain composition has been reported to modulate AKT–membrane interactions, underscoring the functional relevance of PI remodeling beyond headgroup phosphorylation alone.(71) Furthermore, in the context of prostaglandin metabolism and cancer, lack of a regulatory program involving PGD2 synthesis and regulation of PGE2 by HPGD (**Figures 5-7**) fosters the proliferative impact of prostaglandins in non-differentiated colonocytes. Likewise, murine models showed that PTGER4-dependent cell proliferation was involved in colon cancer growth(72), while PTGER2 inhibition led to a significant decrease in cancer initiation(73).

In this study, we propose that a high content in AA-PI species could be involved in sustain colonocyte stemness and that a decrease in content, partly due to prostaglandin metabolism, is involved in the correct differentiation of colonocytes. Our results reinforce Yanes et al.(74) model, in which the membrane unsaturation degree is involved in the required cellular chemical plasticity to maintain a stemness state. Undoubtedly, understanding the role of membrane lipid metabolism in metabolic states defining stem cell capacity for self-renewal and differentiation offers a unique insight to fully comprehend stem cell behavior.(75) Together, these findings define regulatory programs that generate testable mechanistic hypotheses, which will require targeted gain- and loss-of-function studies to establish direct causal relationships.

## CONCLUSIONS

Although the relevance of lipid metabolism is well acknowledged, it has been difficult to advance in the understanding of the regulatory mechanisms. The lack of sensitive and robust technology to run solid lipidomic analysis has lagged the field. Fortunately, the scenario has changed and today, not only is it possible to run thorough high-throughput lipidomic analyses that are fully compatible with clinical practice, but the development of spatially resolved lipidomic methodologies has also firmly established the foundation for integrating high-quality molecular data into anatomy reports. These developments are poised to enhance diagnosis, stratification, and treatment monitoring in Personalized Medicine.

This study provides compelling evidence that membrane lipid composition, particularly the regulation of PI 38:4 and its associated metabolic pathways, plays a pivotal role in colonocyte differentiation and stemness. Through integrative modeling of MSI and transcriptomic data, we identified distinct gene modules that orchestrate lipid remodeling along the crypt axis, revealing a tightly regulated interplay between lipid metabolism and cell fate decisions. The identification of key enzymes and transcriptional regulators, such as PLCD3, PLA2G4F, and HNF4A, underscores the complexity and specificity of lipid-mediated signaling in healthy colonocyte maturation. Moreover, our findings highlight a profound metabolic reprogramming in tumor cells, characterized by sustained PI 38:4 levels and disrupted prostaglandin metabolism, which collectively support aberrant proliferation and impaired differentiation. The loss of the metabolic-like module and the persistence of the proliferative transcriptomic program in cancer further emphasize the pathological significance of lipidome dysregulation. Taken together, these insights reinforce the concept that membrane lipid metabolism is not merely a passive reflection of cellular state but a dynamic and influential determinant of stem cell behavior and tissue homeostasis. Future research targeting lipid metabolic pathways may open new avenues for therapeutic intervention in colorectal cancer and regenerative medicine. The present findings exemplify the analytical depth achievable through spatial lipidomics and demonstrate its potential to generate novel hypotheses addressing complex pathophysiological processes.

## LIST OF ABBREVIATIONS

AA: Arachidonic acid
CM: Co-expression modules
CSC: Colonocyte stem cells
DUFA: Di-unsaturated fatty acid
EPHB2: Ephrin B2 receptor
FACS: Fluorescence-activated cell sorting
kME: Gene-module-membership
MALDI-MSI: Matrix assisted laser desorption/ionization-Mass Spectrometry Imaging
MUFA: Monounsaturated fatty acid
PI: Phosphatidylinositol
PIP: Phosphoinositide
PTG: Prostaglandin
PGE2: Prostaglandin E2
PGF2α: Prostaglandin F2 alpha
PGD2: Prostaglandin D2
PUFA: Polyunsaturated fatty acid
TA cells: Transit-amplifying cells

## RESOURCE AVAILABILITY

Further information and requests for resources and reagents should be directed to and will be fulfilled by lead contact, Gwendolyn Barceló. The gene expression microarray has been deposited to GEO, accession number GSE285090. All data other reported in this paper will be shared by the lead contact upon request. Original code for weighted gene co-expression network analysis and lipid to gene data integration has been deposited at Github, at https://github.com/idisba/lipid-RNA_Maimo-Barcelo, and is publicly available. Additional information upon data reported here is available from the lead contact upon request.

## AUTHOR CONTRIBUTIONS

Conceptualization, A.M-B., J.B-E., J.A.F., R.M.R., and G.B-C.; patient enrolment and sample acquisition, A.M-B., M.A-M., G.P.M, and J.M.O; methodology, A.M-B., J.B-E., K.P-R., L.M.S., M.L.M., D.H.L., C.C., J.A.F.; analysis and visualization, A.M-B., J.B-E., and J.M-F.; writing—original draft, A.M-B., R.M.R, and G.B-C.; writing—review, all authors; supervision, R.M.R. and G.B-C.

## DECLARATION OF INTERESTS

The authors declare that they have no known competing financial interests or personal relationships that could have appeared to influence the work reported in this paper.

## ACKNOWLEDGMENTS

We are greatly thankful to the nurses and medical doctors of the Gastroenterology Department of the Hospital Universitari de Son Espases (HUSE, Palma, Spain) for their participation during patient enrolment and sample acquisition.

## FUNDING

This work was supported in part by the Institute of Health Carlos III (PI19/00002), the Basque Government (IT1491-22) and the EC (European Regional Development Fund, ERDF). English language editing was supported by the Research Committee of the Son Espases University Hospital (HUSE). A. M-B. held a predoctoral contract of the Govern Balear (Direcció General d’Innovació i Recerca) co-funded by ESF (European Social Fund, FPI/2160/2018). J.B-E., held a postdoctoral contract of Health Research Institute of the Balearic Islands, co-funded by Govern de les Illes Balears (ITS 2023 – 057. CONV. FOLIUM-2023-CANCER). K. P-R was supported by a predoctoral contract of the Health Research Institute Carlos III (FI20/00180) co-funded by ESF. L. M-S held a predoctoral fellowship of the Spanish Ministry of Economy, Industry and Competitiveness (BES-2016-078721); R. M-R postdoctoral contract was supported by a grant from the Scientific Foundation of the Spanish Association Against Cancer (INVES222995RODR).

## SUPLEMENTARY MATERIAL

**Supplementary Table 1.** Comprehensive list of the lipid species detected in the epithelial layer of human colon tissue sections with MALDI-MSI (Bestard et al 2016 and Lopez et al 2018). Lipid species are classified according to their distribution patterns along the colon crypt.

**Supplementary Table 2.** Sample clinical data. Healthy and tumor biopsies form colon cancer surgical patients collected and analyzed by MALDI-MSI and gene expression microarrays. M, Male; F, Female; ADC, Adenocarcinoma; G1, well differentiated; G2, moderately differentiated; TNM, Tumor Node Metastasis classification; MMR, Mismatch Repair proteins.

**Supplementary Table 3.** MALDI-MSI lipid data of the isolated colonocyte EPHB2 subpopulations from healthy and tumor human samples. Excel file includes figure 1 data: RAW data, lipid assignation, Mole species normalization, PI and PE MUFA-PUFA data (excel file).

**Supplementary Table 4.** GSEA TOP 20 ranked human MSigDB collection hallmark gene sets (H: 55 gene sets) and curated gene sets (C2:Reactome 1654 gene sets21) enriched pathways in EPHB2 High vs. Low gene expression samples. Excel file includes detailed summary of enrichment analysis.

**Supplementary Table 5.** List of genes related to lipid metabolism, according to GO, REACTOME and Wikipathways databases (Txt file).

**Supplementary Table 6.** Top 20 enriched lipid pathways of healthy vs. tumor gene expression samples, and in EPHB2 High vs. Low gene expression analysis. Pathway enrichment was calculated with TAC software (Thermo Fisher Scientific) and using Wikipathways database. Excel file includes detailed results of the enrichment analysis(excel file).

**Supplementary Table 7.** Multiomics integration matrix and GO enrichment of WGCNA gene modules. WGCNA gene composition of PI 38:4 and PI 36:1 modules (excel file).

**Supplementary Table 8.** a) Network lipid gene composition and reactome enrichment. b) Gene module membership and gene significance for PI 38:4 in colonocyte differentiation (excel file).

**Supplementary Table 9.** Summary of gene modules top ranked gene drivers (WGCNA) and significantly enriched transcription factors (ChIP-Seq TF binding sites, iCisTarget). Healthy (H), Tumor (T). Extended data of iCisTarget results on dedicated supplementary files (ZIP file).

**Supplementary Figure 1.** Unsupervised hierarchical clustering heatmap of differential gene expression analysis of Healthy vs. Tumor samples.

**Supplementary Figure 2.** The differential expression analysis of Healthy (a) and tumor High-EPHB2 vs. Low-EPHB2.

**Supplementary Figure 3.** Gene Set Enrichment Analysis (GSEA) for cell-type-specific gene signatures from the large intestine in the human Molecular Signatures Database (MSigDB)19 in EPHB2High vs. Low.

**Supplementary Figure 4.** Changes in organoid size during PLA2 and PTGS inhibition. Representative images of the organoids cultured at different PLA2 and PTGS/COX inhibitors concentration.

**Supplementary Figure 5.** Assessment of the PLA2 and PTGS inhibition capacity of the compounds used.

**Supplementary Figure 6.** Changes in organoid size and gene expression subtype markers during PLA2 and PTGS inhibition. Effects of PLA2 + PTGS inhibition over mice colon organoid proliferation and differentiation.

**Supplementary Figure 7.** Mander’s coefficient value of EP1 (PTGER1) receptor along crypt colonocytes nuclei (base and middle to luminal side).

**Supplementary Figure 8.** Representative immunofluorescence assay of each prostaglandin receptor tested and the ATPase Na+/K+ as a negative control of nuclear presence.

**Supplementary Figure 9.** MTTassay of pharmacological inhibitors and prostaglandin treatment in colon organoids.

**Supplementary Figure 10.** Effect on organoid growth rate at 48h upon PTGER4 agonist and antagonist, PTGS and PLA2 inhibition and prostaglandin treatments.

**Supplementary Figure 11.** Tuft cell markers DCLK1, HPGDS, PTGS1, and KIT gene expression decreases in colon cancer samples compared to normal tissue.

**Supplementary Figure 12.** Representative configuration of FACSAria Fusion with FACSDiva v8.0 (BD Biosciences) gating for EPHB2 subpopulations.

**Supplementary Figure 13.** Weighted Gene Co-expression Network Analysis.

## Notes

### Competing Interest Statement

The authors have declared no competing interest.

