## Supplementary material and data tables for "“A Fine-Tuned Phosphatidylinositol Profile Contributes to Colonocyte Differentiation and Malignization: Evidence From Integrated Omics”": Supplementary material Maimo-Barcelo et al_Rvwd.docx

### Supplementary material: “Phosphatidylinositol Remodeling Marks the Loss of Differentiation in Colon Cancer”

#### **ACKNOWLEDGMENTS**

We are greatly thankful to the nurses and medical doctors of the Gastroenterology Department of the Hospital Universitari de Son Espases (HUSE, Palma, Spain) for their participation during patient enrolment and sample acquisition.

#### **FUNDING**

This work was supported in part by the Institute of Health Carlos III (PI19/00002), the Basque Government (IT1491-22) and the EC (European Regional Development Fund, ERDF). English language editing was supported by the Research Committee of the Son Espases University Hospital (HUSE). A. M-B. held a predoctoral contract of the Govern Balear (Direcció General d'Innovació i Recerca) co-funded by ESF (European Social Fund, FPI/2160/2018). J.B-E., held a postdoctoral contract of Health Research Institute of the Balearic Islands, co-funded by Govern de les Illes Balears (ITS 2023 – 057. CONV. FOLIUM-2023-CANCER). K. P-R was supported by a predoctoral contract of the Health Research Institute Carlos III (FI20/00180) co-funded by ESF. L. M-S held a predoctoral fellowship of the Spanish Ministry of Economy, Industry and Compactivity (BES-2016-078721); R. M-R postdoctoral contract was supported by a grant from the Scientific Foundation of the Spanish Association Against Cancer (INVES222995RODR).

#### **Supplementary Table 1.** Comprehensive list of the lipid species detected in the epithelial layer of human colon tissue sections with MALDI-IMS (Bestard et al 2016 and Lopez et al 2018). Lipid species are classified according to their distribution patterns along the colon crypt.

| Gradual distribution (nº of detected species = 27) | | | | | | |
| --- | --- | --- | --- | --- | --- | --- |
| Lipid class | **Assigned lipid species** | **R^2^** | **Adjustment mode^(a)^** | | **Gradient^(b)^** | **RI basal / apical values^(c)^** |
| PI | | | | | | |
|  | 32:1 | 0.62 | Log | | + | 0.2/0.6 |
|  | 34:1 | 0.89 | lineal | | + | 8.0/17.0 |
|  | 34:2 | 0.76 | Log | | + | 5.0/8.0 |
|  | 36:1 | 0.98 | lineal | | + | 6.0/20.0 |
|  | 36:2 | 0.94 | Log | | + | 5.7/27.0 |
|  | 36:3 | 0.76 | 2^nd^ | | - |  |
|  | 36:4 | 0.94 | Log | | - | 8.2/2.5 |
|  | 38:4 | 0.97 | Log | | - | 71.6/15.0 |
|  | 38:5 | 0.81 | lineal | | - | 2.5/1.5 |
|  | 40:4 | 0.82 | lineal | | - | 1.4/0.4 |
| PE | | | | | | |
|  | 34:1 | 0.80 | Log | | + | 4.3/7.2 |
|  | 36:3 | 0.85 | 2^nd^ | | - |  |
|  | 36:4 | 0.42 | lineal | | - | 1.8/1.0 |
|  | 38:1 | 0.56 | 2^nd^ | | + |  |
|  | 38:4 | 0.79 | Log | | - | 20.0/5.4 |
|  | 38:5 | 0.40 | lineal | | - | 3.6/1.6 |
| PC | | | | | | |
|  | 32:0 | 0.69 | lineal | | - | 6.5/4.6 |
|  | 38:2 | 0.68 | lineal | | - | 0.7/0.4 |
|  | 38:3 | 0.68 | lineal | | - | 1.0/0.5 |
|  | 38:4 | 0.43 | Log | | - | 2.8/1.1 |
|  | 38:5 | 0.26 | Log | | - | 1.1/0.7 |
| PE P | | | | | | |
|  | 16:0/18:1 | 0.70 | Log | | + | 8.3/14.4 |
|  | 18:0/18:1, (PC (P-16:0/18:1)) | 0.82 | Log | | + | 5.6/13.0 |
|  | 18:0/18:2, 18:1/18:1, (PC-P(16:0/18:2)) | 0.63 | Log | | + | 8.3/15.5 |
|  | 16:0/20:4 | 0.91 | Log | | - | 21.3/9.7 |
|  | 18:0/20:4, 16:0/22:4 | 0.77 | Log | | - | 28.6/16.1 |
|  | 20:0/20:4, 18:0/22:4, (PC (P-18:0/20:4)) | 0.84 | Log | | - | 8.1/1.9 |
| Even distribution (nº of detected species = 15) | | | | | | |
| Lipid class | **Assigned lipid species** | **Mean** | | **SD** | | |
| PE | | | | | | |
|  | 36:1 | 24.23 | | 1.34 | | |
|  | 36:2 | 25.41 | | 1.97 | | |
|  | 38:2 | 9.99 | | 0.91 | | |
|  | 38:3 | 7.29 | | 0.70 | | |
| PC | | | | | | |
|  | 32:1 | 1.50 | | 0.11 | | |
|  | 34:1 | 37.17 | | 1.52 | | |
|  | 34:2 | 16.40 | | 0.62 | | |
|  | 34:3 | 1.50 | | 0.10 | | |
|  | 36:2 | 17.10 | | 0.40 | | |
|  | 36:3 | 10.40 | | 0.74 | | |
| PE P | | | | | | |
|  | 16:0/18:2 | 8.4 | | 0.9 | | |
|  | 16:0/20:3, 18:1/18:2 | 4.8 | | 0.6 | | |
|  | 18:1/20:4, 16:0/22:5, 18:0/20:5 | 8.8 | | 1.5 | | |
|  | 16:0/22:6 | 4.4 | | 0.6 | | |
|  | 18:0/22:6, 18:1/22:5, (PC(P-16:0/22:6)) | 4.7 | | 0.6 | | |
| Scattered distribution (nº of detected species = 5) | | | | | | |
| Lipid class | Assigned lipid species | mean | | SD | | |
| PI | | | | | | |
|  | 32:0 | 0.53 | | 0.06 | | |
|  | 38:6 | 0.99 | | 0.20 | | |
| PE | | | | | | |
|  | 34:2 | 2.27 | | 0.31 | | |
| PE | | | | | | |
|  | 36:1 | 4.44 | | 0.49 | | |
|  | 36:4 | 3.20 | | 0.43 | | |

^(a)^ Variations in relative intensity (RI) along the crypt were modeled using either a linear equation, a second-degree polynomial, or a logarithmic function. ^(b)^ The slope was defined as positive when RI increased from the crypt base toward the lumen, and negative when it decreased in that direction. ^(c)^ To quantify the extent of change, RI values at the initial and final pixels, representing the base and top of the crypt, were reported. These values are shown as percentages of the total RI for a specific lipid class and reflect the average RI from five separate crypts measured across four independent experiments.

#### **Supplementary Table 2.** Sample clinical data. Healthy and tumor biopsies form colon cancer surgical patients collected and analyzed by MALDI-MSI and gene expression microarrays. M, Male; F, Female; ADC, Adenocarcinoma; G1, well differentiated; G2, moderately differentiated; TNM, Tumor Node Metastasis classification; MMR, Mismatch Repair proteins.

| **Patient** | **Age** | **Sex** | **Localization** | **Histological type** | **Tumor Differentiation. Grade** | **TNM** | **MMR IHQ** | **Omic analysis** |
| --- | --- | --- | --- | --- | --- | --- | --- | --- |
| 1 | 54 | M | Sigmoid colon | ADC | G2 | pT3, N2a, M0 | MLH1, PMS2, MSH2, MSH6 (+) | Healthy cells MALDI-MSI |
| 2 | 65 | M | Sigmoid colon | ADC | G2 | pT4b, N0, M0 | MLH1, PMS2, MSH2, MSH6 (+) | Healthy cells MALDI-MSI |
| 3 | 83 | F | Sigmoid colon | ADC | G1 | T3, N0, M0 | MLH1, PMS2, MSH2, MSH6 (+) | Healthy cells MALDI-MSI |
| 4 | 75 | F | Sigmoid colon | ADC | G2 | T3, N1c, M0 | MLH1, PMS2, MSH2, MSH6 (+) | Healthy cells MALDI-MSI |
| 5 | 71 | M | Ascending colon | ADC | G2 | T3, N0, M0 | MLH1, PMS2, MSH2, MSH6 (+) | Tumor cells MALDI-MSI |
| 6 | 86 | M | Ascending colon | ADC | G2 | T3, N0, M0 | MLH1, PMS2, MSH2, MSH6 (+) | Tumor cells MALDI-MSI |
| 7 | 71 | F | Ascending colon | ADC | G2 | T3, N1c, M0 | MLH1, PMS2, MSH2, MSH6 (+) | Tumor cells MALDI-MSI |
| 8 | 82 | M | Sigmoid colon | ADC | G2 | T3, N2a, M0 | MLH1, PMS2, MSH2, MSH6 (+) | Tumor cells MALDI-MSI |
| 9 | 48 | M | Sigmoid colon | Mucinous ADC | G1 | T4b, N0, M0 | MLH1, PMS2, MSH2, MSH6 (+) | Tumor cells MALDI-MSI |
| 10 | 65 | M | Sigmoid colon | ADC | G2 | T3, N0, M0 | MLH1, PMS2, MSH2, MSH6 (+) | Healthy and Tumor cells MALDI-MSI |
| 11 | 85 | F | Sigmoid colon | ADC | G1 | T3, N0, M0 | MLH1, PMS2, MSH2, MSH6 (+) | Healthy and Tumor cells Human clariom S pico Affymetrix |
| 12 | 72 | F | Ascending colon | ADC | G2 | T4a,N0, M0 | MLH1(-) PMS2(-) MSH2(+) MSH6 (+) | Healthy cells MALDI-MSI |
| 13 | 71 | M | Sigmoid colon | ADC | G2 | T3, N2a, M0 | MLH1, PMS2, MSH2, MSH6 (+) | Healthy cells MALDI-IMS |
| 14 | 68 | F | Ascending colon | Mucinous ADC | G2 | T3,N0, M0 | MLH1(-), PMS2(-), MSH2(+), MSH6 (+) | Healthy and Tumor cells Human clariom S pico Affymetrix |
| 15 | 78 | M | Ascending colon | ADC | G2 | T3,Nb, M0 | MLH1, PMS2, MSH2, MSH6 (+) | Healthy and Tumor cells Human clariom S pico Affymetrix |
| 16 | 83 | M | Ascending colon | ADC | G1 | T3, N0, M0 | MLH1, PMS2, MSH2, MSH6 (+) | Healthy and Tumor cells Human clariom S pico Affymetrix |
| 17 | 55 | M | Sigmoid colon | ADC | G2 | pT3,N1a,M0 | MLH1, PMS2, MSH2, MSH6 (+) | Healthy cells MALDI-IMS |

#### **Supplementary Table 3.** MALDI-IMS lipid data of the isolated colonocyte EPHB2 subpopulations from healthy and tumor human samples. Excel file includes figure 1 data: RAW data, lipid assignation, Mole species normalization, PI and PE MUFA-PUFA data.

#### **Supplementary Table 4.** GSEA TOP 20 ranked human MSigDB collection hallmark gene sets (H: 55 gene sets) and curated gene sets (C2:Reactome 1654 gene sets21) enriched pathways in EPHB2 High *vs*. Low gene expression samples. Excel file includes detailed summary of enrichment analysis.

#### **Supplementary Table 5.** List of genes related to lipid metabolism, according to GO, REACTOME and Wikipathways databases. Txt file.

#### **Supplementary Table 6.** Top 20 enriched lipid pathways of healthy vs. tumor gene expression samples, and in EPHB2 High *vs*. Low gene expression analysis. Pathway enrichment was calculated with TAC software (Thermo Fisher Scientific) and using Wikipathways database. Excel file includes detailed results of the enrichment analysis.

#### **Supplementary Table 7:** a) Multiomics integration matrix and GO enrichment of WGCNA gene modules. b) WGCNA gene composition of PI 38:4 and PI 36:1 modules (figure 5). Excel file includes the integration data and gene composition of CMs.

#### **Supplementary Table 8:** a) Network lipid gene composition and reactome enrichment. b) Gene module membership and gene significance for PI 38:4 in colonocyte differentiation. Excel file contains all the data.

#### **Supplementary Table 9.** Summary of gene modules top ranked gene drivers (WGCNA) and significantly enriched transcription factors (ChIP-Seq TF binding sites, iCisTarget). Healthy (H), Tumor (T). Extended data of iCisTarget results on dedicated supplementary files.

| **Sample** | **Module** | **Trait** | **CM-trait Correlation** | **Top-ranked CM Gene drivers** | **Top-ranked Transcription factors** |
| --- | --- | --- | --- | --- | --- |
| H | Turquoise | PI 38:4 | + | *CHRNAS, SLC26A3, CA7, CA1, HRCT1* | (-)kME: SIN3A (NES 5.41), JUND (AP-1, NES 6.21), and EVI1 (NES 5.24)  (+)kME: E2F4 (NES 11.58), TFDP1 (NES 10.88), and E2F7 (NES 8.69) |
| H | Brown | PI 38:4 | - | *COL17A1, TRIM40, OXGR1, MORC4, RAD18* | (-)kME: ETV5(ERM, NES 9.15), HNF4A (NES 5.68), EP300 (NES 4.22)  (+)kME: MBD4 (NES 7.3), NFIC (NES 5.28), NFE2L2 (NES 4.13) |
| H | Red | PI 36:1 | - | *SMG8, FER, SPEN, SMURF2, OR10K2* | (-)kME: EZH2 (NES 6.41), CBX2 (NES 6.23) and CBX8 (NES 5.69)  (+)kME: PHF8 (NES 5.31), TAF1 (NES 4.57), and MYC (NES 3.71) |
| T | Dark magenta | PI 38:4 | + | *TOX2, ZNF317, CDKAL1, LINC00663, HS2ST1* | (-)kME: CBX8 (NES 9.80), CEBPB (NES 5.81), and ELF1 (NES 5.29)  (+)kME: REST (NES 5.65) , EP300 (NES 4.82), and PCR1X (NES 4.68) |
|  |  | PI 36:1 | - | *TOX2, ZNF317, LINC00663, AICDA, RNF144B* |  |
| T | Dark red | PI 36:1 | + | *ZNF200, C1orf210, THSD4, CDR2, AKR1C1* | (-)kME: HNF4G (NES 6.97), TCF12 (NES 6.96), and NR2F2 (NES 6.84)  (+)kME: TFDP1 (NES 25.91), E2F4 (NES 20.07), and HDAC1 (NES 13.04) |
|  |  | PI 38:4 | - | *MNS1, GRAMD3, ATAD2, BARD1, CYP27A1* |  |
| T | Turquoise | EPHB2 | + | *SRP68, NUP160, MSH2, TRA2B, RECQL4* | (-)kME: MBD4 (NES 4.53), POLR2A (NES 4.01), and EZH2 (NES 3.909  (+)kME: E2F4 (NES 9.86), TFDP1 (NES 9.87), and E2F7 (NES 6.87) |
| T | Green | EPHB2 | + | *DUSP3, ZNF846, DPY19L2, CHRFAM7A, RFX3* | (-)kME: POLR2A (NES 12.75), RAD21 (NES 3.98), and SMC3 (3.85)  (+)kME: CBX2 (NES 4.23), EZH2 (NES 4.23), and GATA3 (NES 4.10) |

#### **Supplementary Figure 1:** Unsupervised hierarchical clustering heatmap of differential gene expression analysis of Healthy vs. Tumor samples. Gene expression data were filtered by false discovery rate (FDR ≤ 0.05) and fold change (FC, ≤ -4 or ≥ 4) resulting in 343 differentially expressed genes, 161 were tumor up-regulated, and 182 down-regulated (left). **b)** Data filtering based on lipid-related genes (Suppl. Table 5), shows 169 differentially expressed genes, 62 up- and 107 down-regulated genes in tumor subpopulations (FDR, ≤ 0.05, FC ≤ -2 or ≥ 2) (right). **c)** Gene expression of colonocyte stem cell markers EPHB2, LGR5, OLFM4; cell proliferation marker MKI67, and mature enterocyte KRT20 in both healthy and tumor isolated EPHB2 colonocytes.

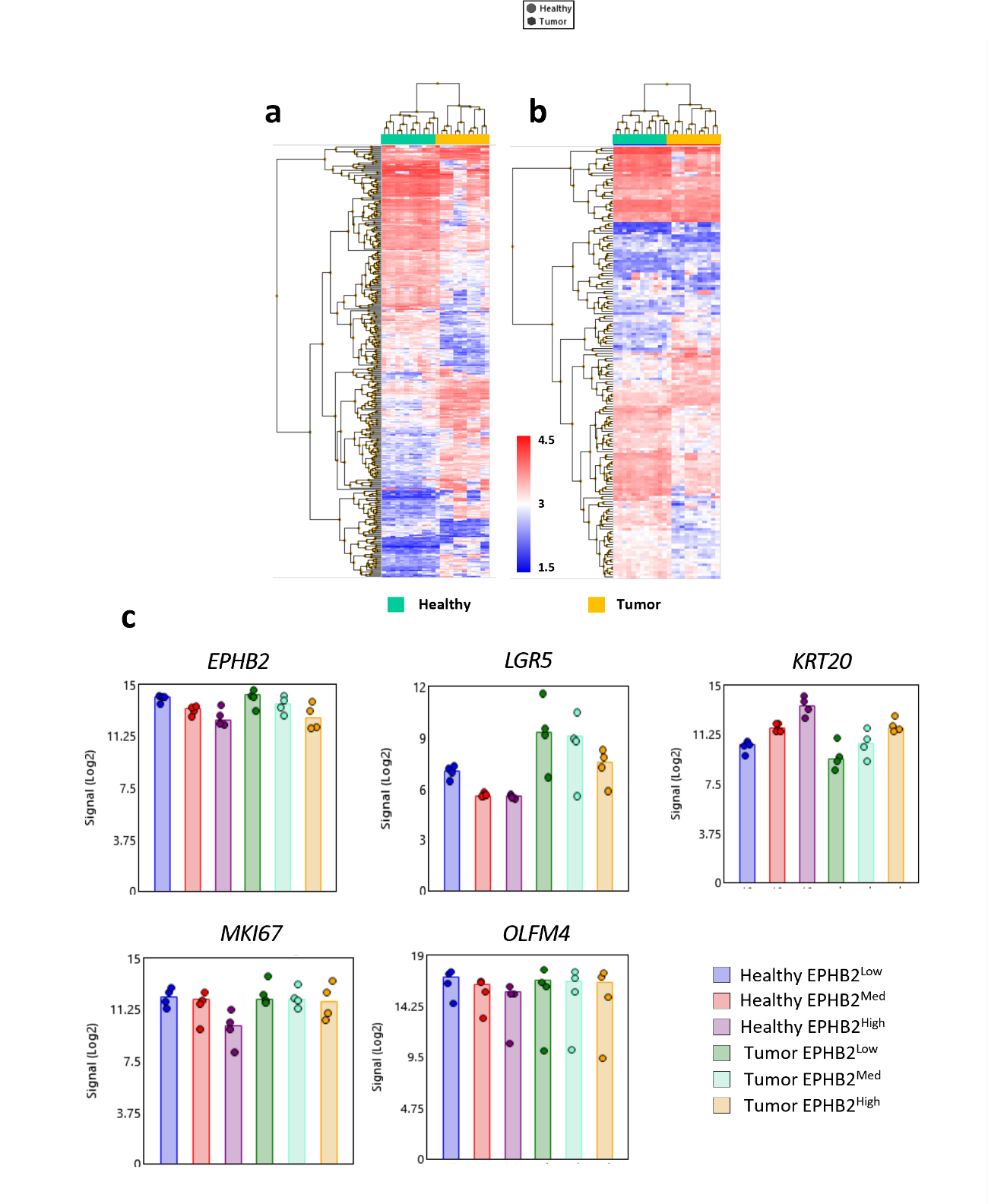

#### **Supplementary Figure 2.** The differential expression analysis of Healthy (a) and tumor High-EPHB2 vs. Low-EPHB2. Gene-Level Fold Change < -2 or > 2, P-Value < 0.05, EbayesAnova Method. A Probeset (Gene/Exon) is considered expressed if ≥ 50% samples have DABG values below DABG Threshold, DABG < 0.05.

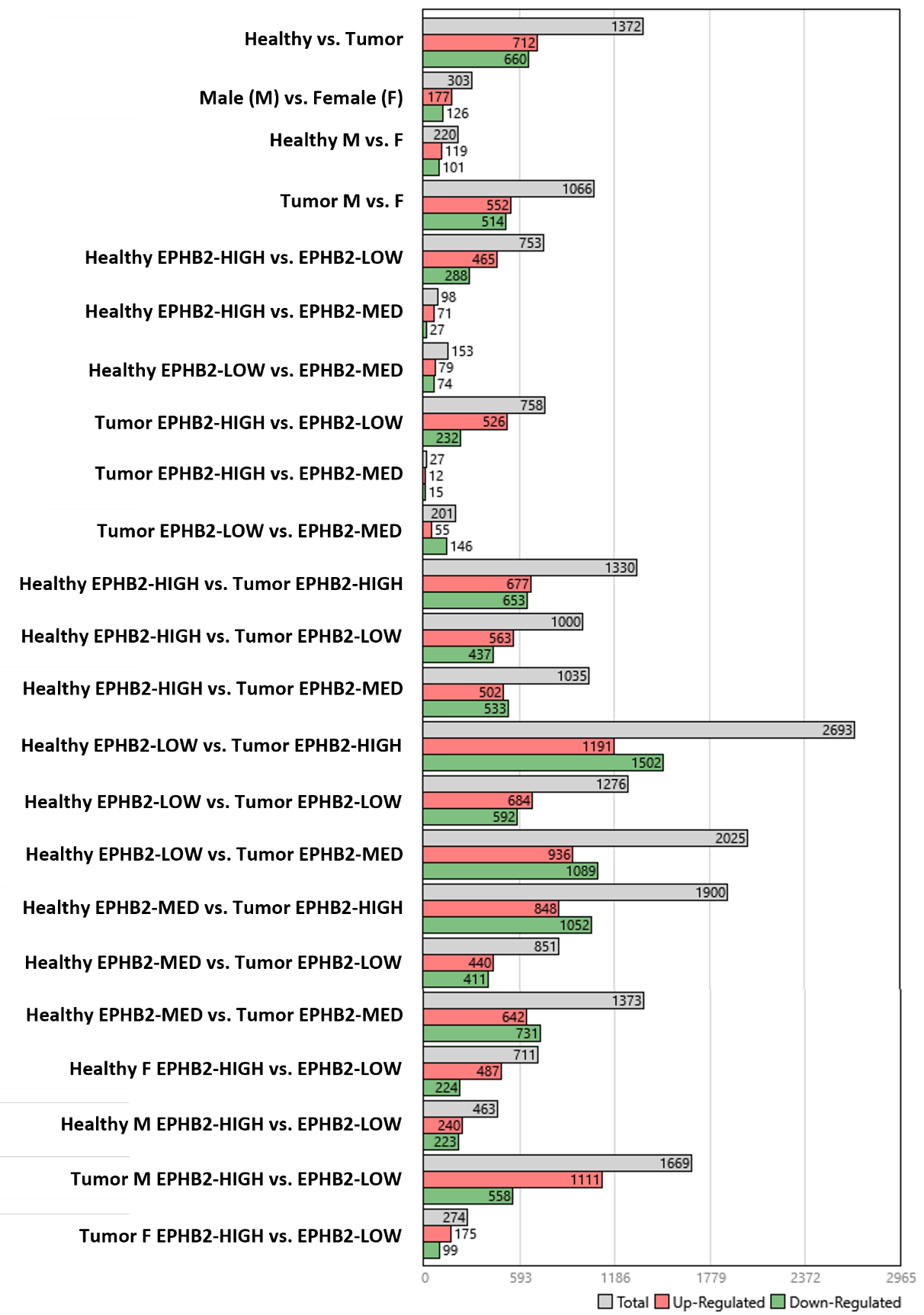

#### **Supplementary Figure 3.** Gene Set Enrichment Analysis (GSEA) for cell-type-specific gene signatures from the large intestine in the human Molecular Signatures Database (MSigDB)^19^ in EPHB2^High^ *^vs^*^. Low^.

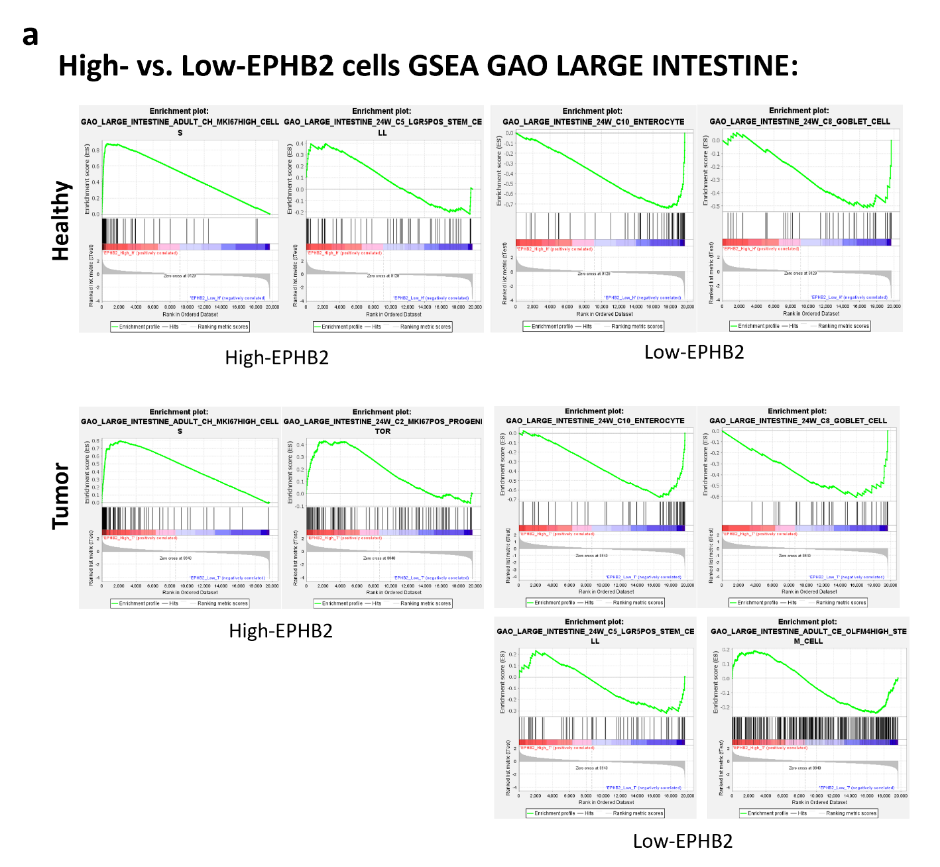

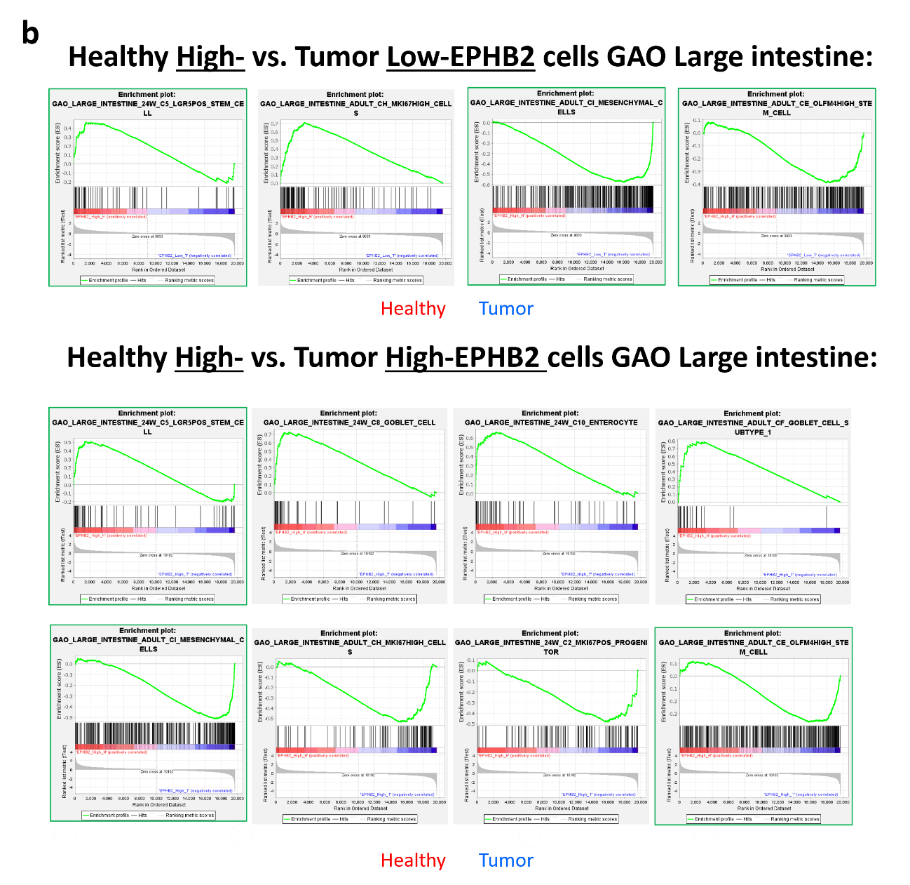

#### **Supplementary Figure 4.** Changes in organoid size during PLA2 and PTGS inhibition. Representative images of the organoids cultured at different PLA2 and PTGS/COX inhibitors concentration. The maximum concentration of each inhibitor before organoid development impairment compared to control conditions was: 5 μM for Arachidonyl trifluoromethyl ketone (ATK), 20 μM for Bromoenol lactone (BEL), 0.1 mM for Valeroyl salicylate (VS) and 4 μM for Celecoxib (Cel). The increase in the organoid size is a reflection of their proliferative capacity; therefore, cultured organoids size was first evaluated by measuring their main area upon 48h.

| **Vehicle (DMSO 0.3%)** | | | | |
| --- | --- | --- | --- | --- |
| 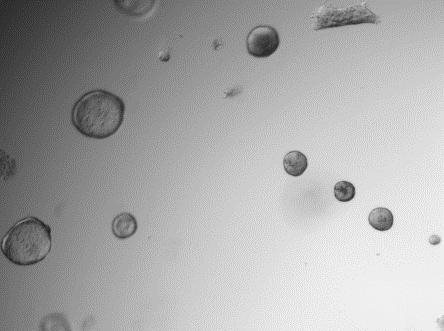 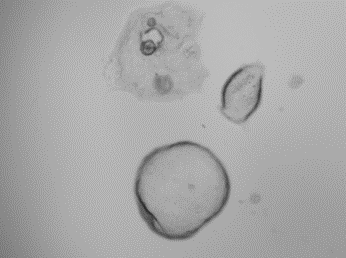 | | | | |
| **Arachidonyl trifluoromethyl ketone (ATK)** | | | | |
| 1 μM  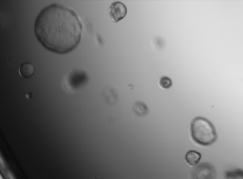 | 2 μM  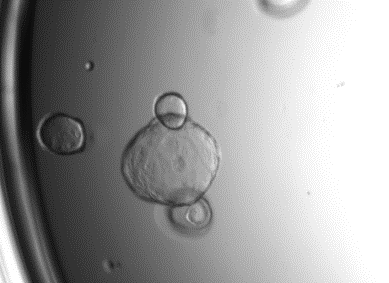 | 5 μM  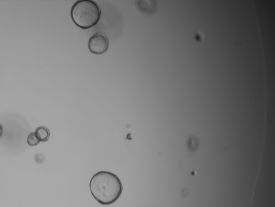 | 10 μM  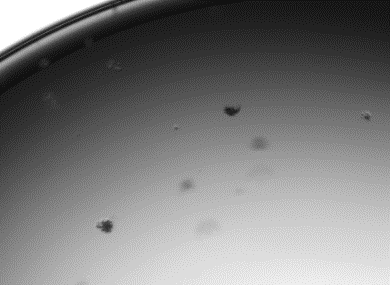 | 20 μM  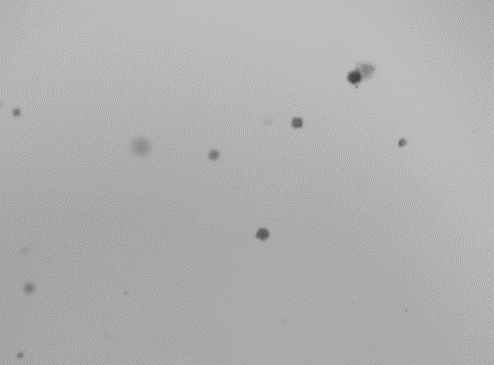 |
| **Bromoenol lactone (BEL)** | | | | |
| 1 μM  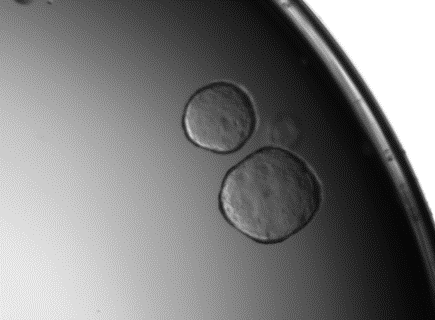 | 2 μM  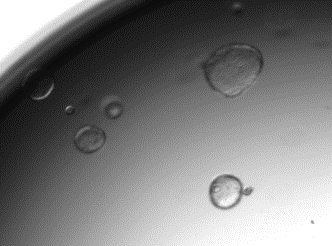 | 20 μM  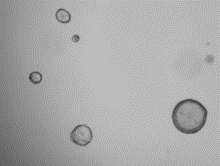 | 50 μM  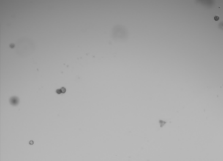 | 100 μM  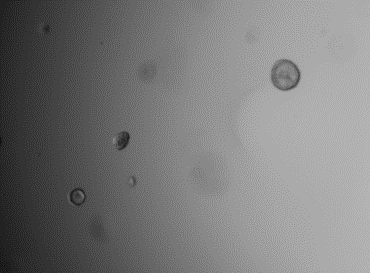 |
| **Valeroyl salicilate (VS)** | | | | |
| 0.05 mM  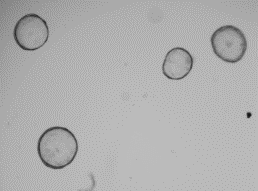 | 0.1 mM  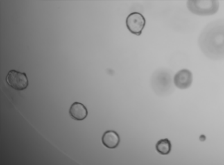 | 0.2 mM  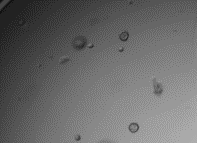 | 0.5 mM  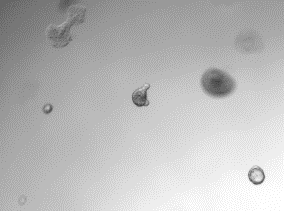 | 0.8 mM  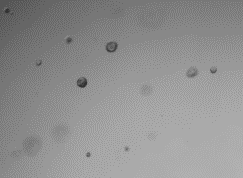 |
| **Celecoxib (CEL)** | | | | |
| 2 μM  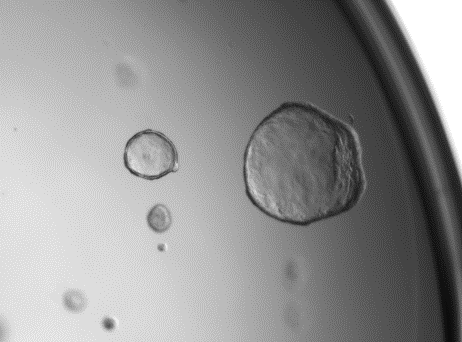 | 4 μM  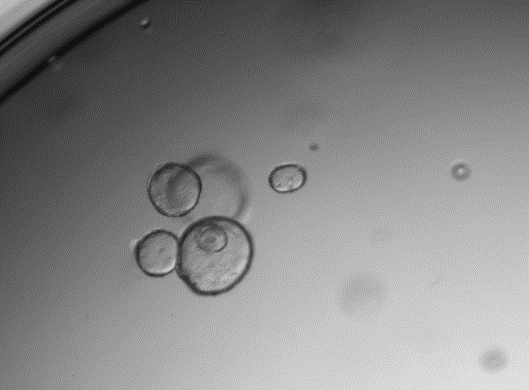 | 6 μM  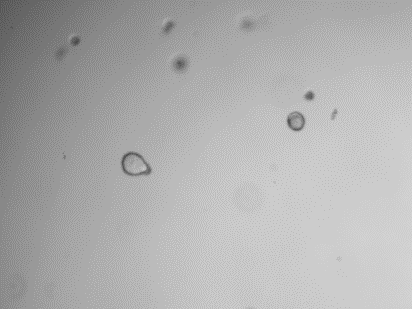 | 10 μM  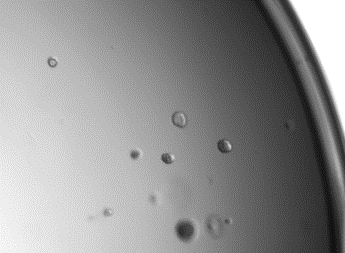 | 20 μM  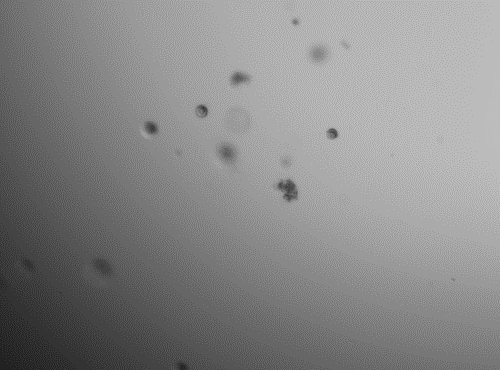 |

#### **Supplementary Figure 5.** Assessment of the PLA2 and PTGS inhibition capacity of the compounds used. PLA2 activity after treatment with PLA2 inhibitors using the EnzChek Phospholipase A2 assay kit of Invitrogen. The concentrations used for each inhibitor were: 5 μM for ATK and 20 μM for BEL (left-side diagram). PTGS activity after treatment with inhibitors using the COX Activity Assay kit (Abcam) (right-side diagram). The concentrations used were: 0.1 mM for VS and 4 μM for Cel. DMSO: dimethylsulfoxyde (DMSO), arachidonyl trifluoromethyl ketone (ATK), bromoenol lactone (BEL), combination of ATK and BEL (PLA2), valerolyl salycilate (VS), celecoxib (Cel), combination of VS and Cel (COX). Values represent mean±SEM (n=4-8). Statistical significance was assessed using ANOVA followed by Bonferroni post-test analysis. *** P<0.001.

| 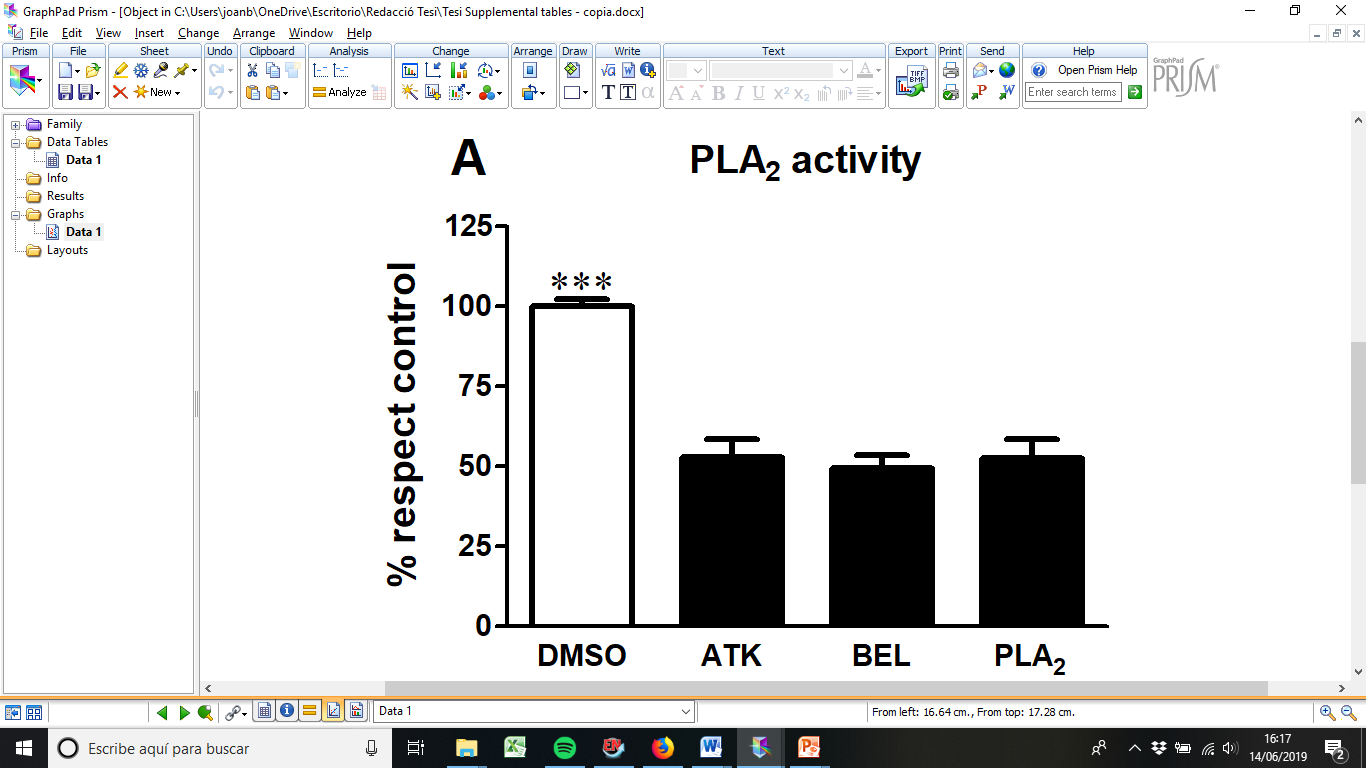 | 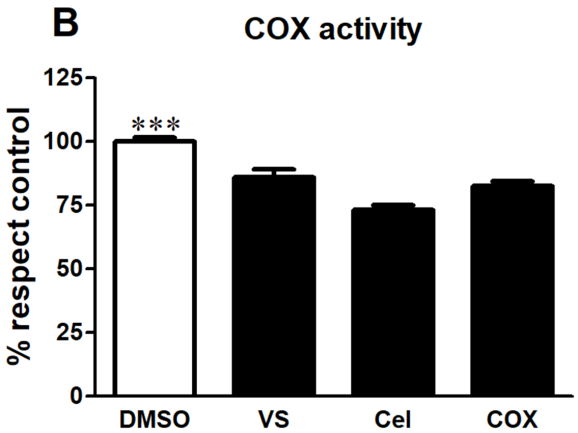 |
| --- | --- |

#### **Supplementary Figure 6.** Changes in organoid size and gene expression subtype markers during PLA2 and PTGS inhibition. Effects of PLA2 + PTGS inhibition over mice colon organoid proliferation and differentiation. A) Colon organoid growth. The growth ratio was calculated by dividing the well value at 48h by the mean value at 0h. The experiment was carried out in 8 different animals. Statistical differences were established using one-way ANOVA followed by Bonferroni post-test. B-C) Representative images of organoids for each treatment B) organoids treated with vehicle, C) organoids treated with PLA2 and PTGS inhibitors, scale bar= 50 µm. D-G) Colon organoid populations assessed by ddPCR. D) Wdr43 expression, E) cKit expression, F) Lgr5 expression G) Krt20 expression. ddPCR values are referred to control and represent the mean ± SEM, n=3-7. To assess statistical differences t-test analysis was applied. Inhibitor concentrations: 5 μM for Arachidonyl trifluoromethyl ketone (ATK), 20 μM for Bromoenol lactone (BEL), 0.1 mM for Valeroyl salicylate (VS) and 4 μM for Celecoxib (Cel).

| **A**  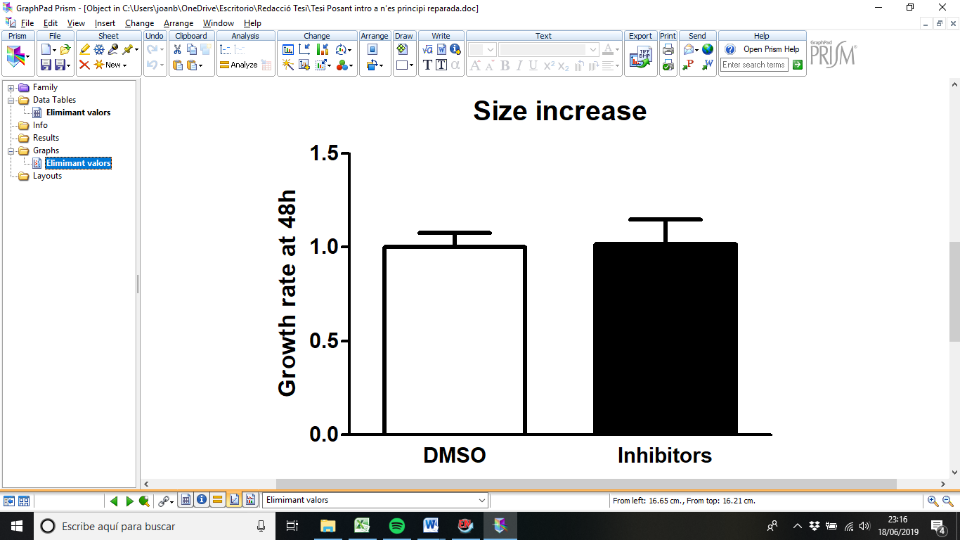 | 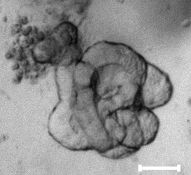  **B**  **C** |
| --- | --- |
|  | 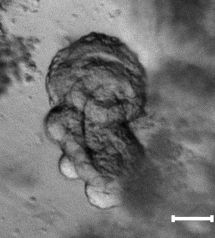 |
| 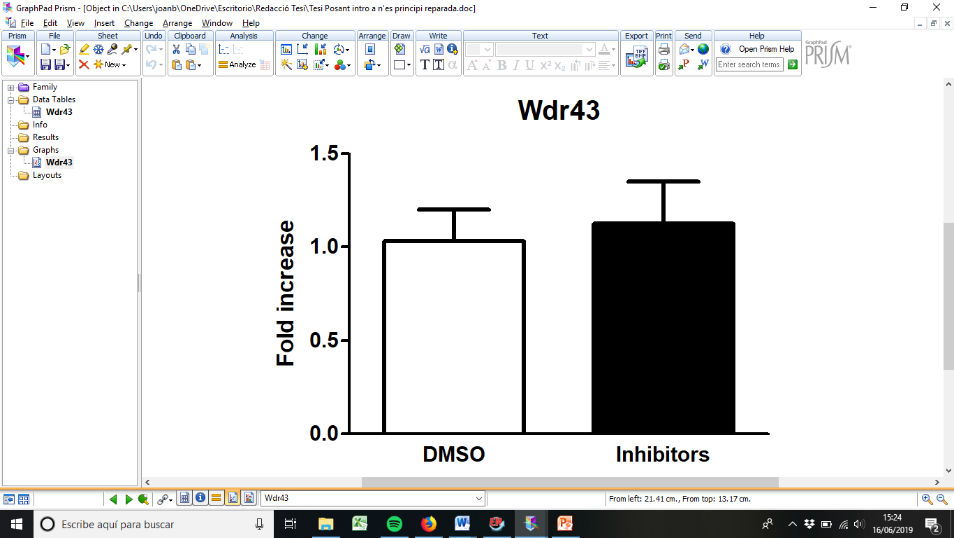  **E**  **D** | 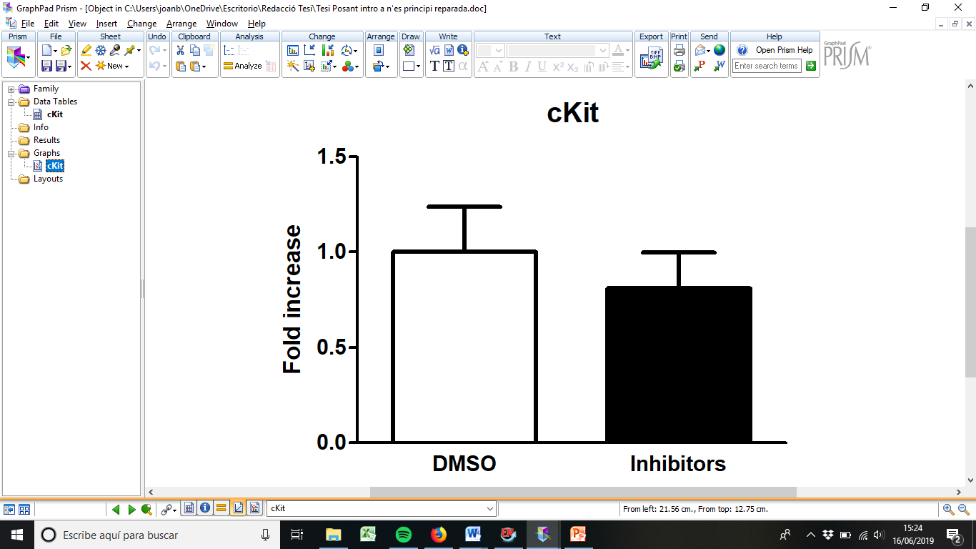 |
| 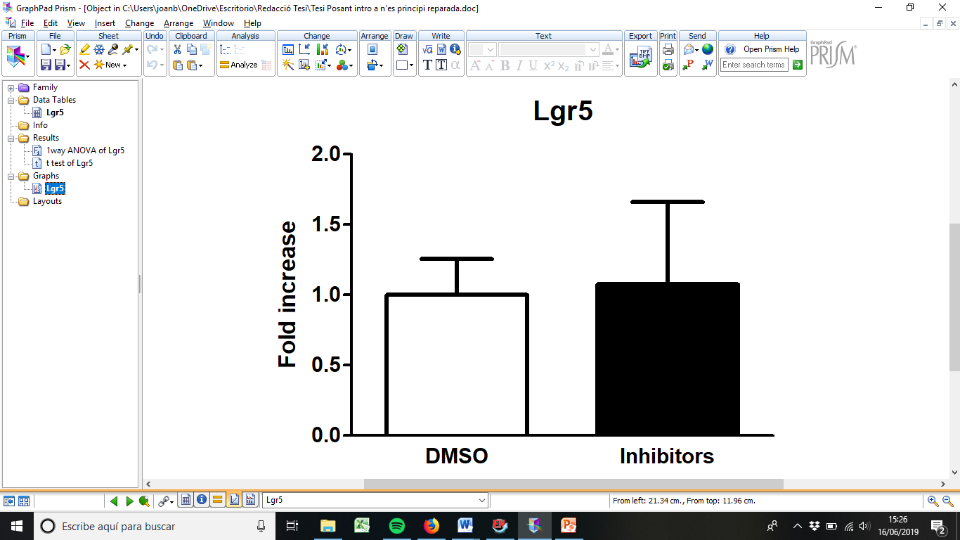  **G**  **F** | 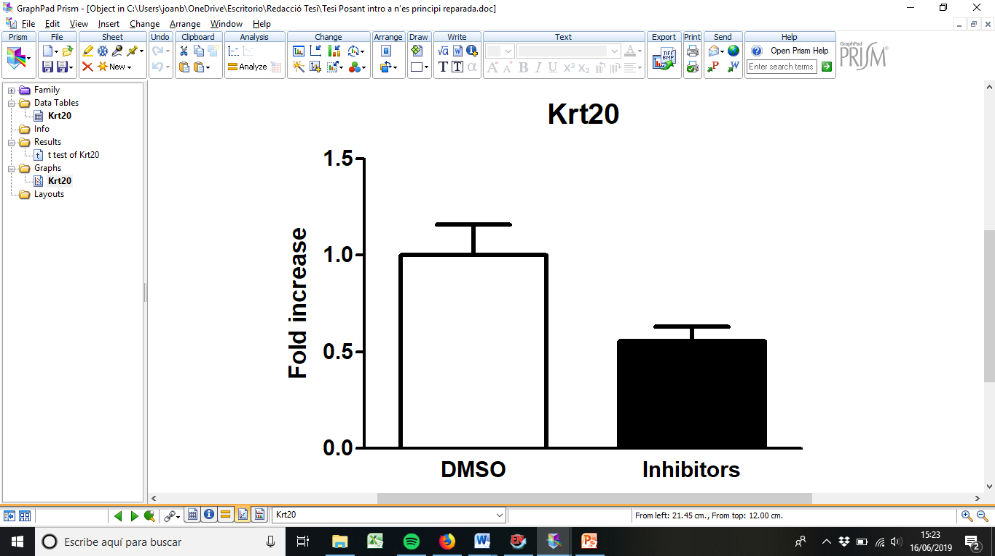 |

#### **Supplementary Figure 7.** Mander’s coefficient value of EP1 (PTGER1) receptor along crypt colonocytes nuclei (base and middle to luminal side). Immunofluorescence experiments were made on NMRI mouse colon, ATPase Na+, K+ was used as negative control and DAPI for nuclei staining. Values represent mean ± SEM (10nuclei/crypt, 5 crypts; 3 consecutive sections; for n=3 mouse). Statistical difference was assessed with t-test analysis, ** P<0.001.

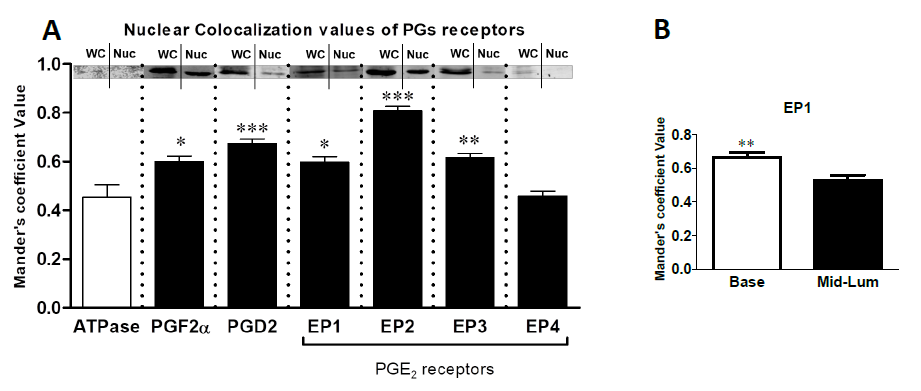

#### **Supplementary Figure 8.** Representative immunofluorescence assay of each prostaglandin receptor tested and the ATPase Na+/K+ as a negative control of nuclear presence.

|  | **DAPI** | **Antibody** | **Merge** |
| --- | --- | --- | --- |
| **ATPase Na^+^/K^+^** | 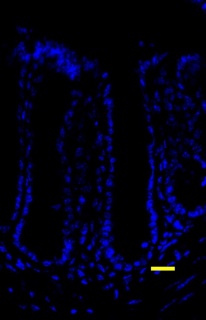 | 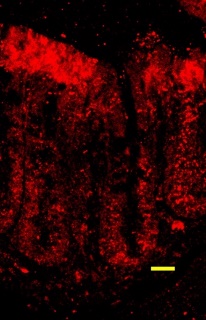 | 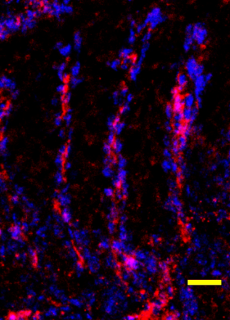 |
| **PGF_2α_ receptor** | 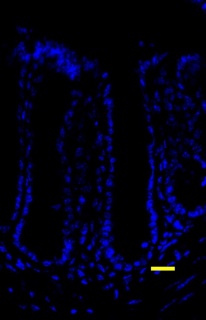 | 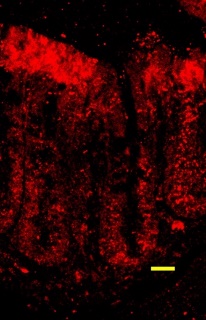 | 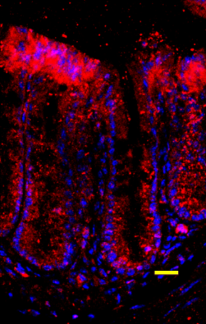 |
| **PGD_2_ receptor** | 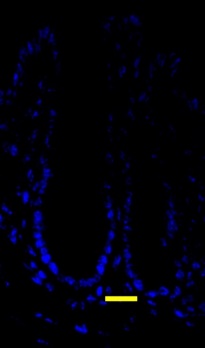 | 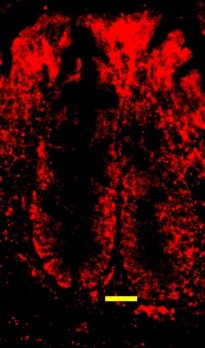 | 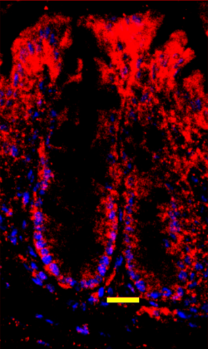 |
| **EP1** | 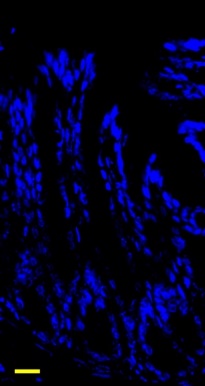 | 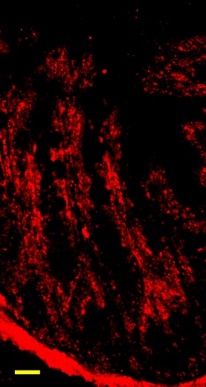 | 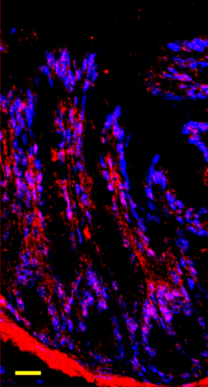 |
| **EP2** | 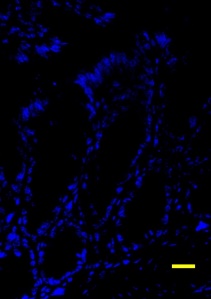 | 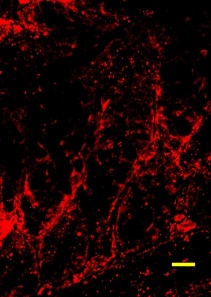 | 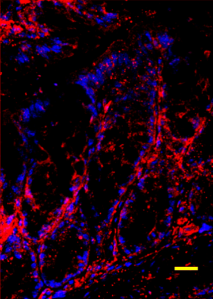 |
| **EP3** | 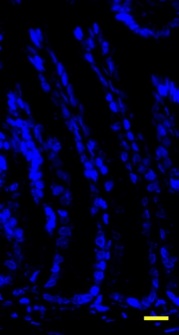 | 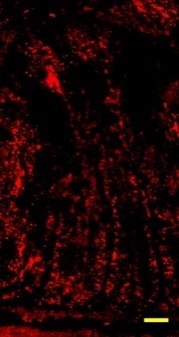 | 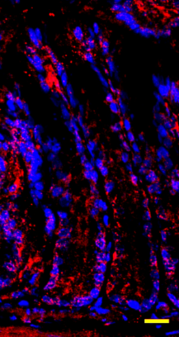 |
| **EP4** | 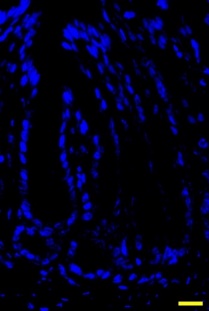 | 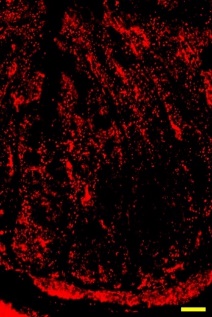 | 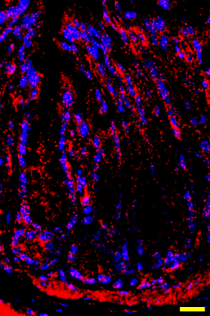 |

#### **Supplementary Figure 9.** MTTassay of pharmacological inhibitors and prostaglandin treatment in colon organoids. **a)** MTT assay was used to test the toxicity of: 5 μM for Arachidonyl trifluoromethyl ketone (cPLA2 inhibitor); 20 μM for Bromoenol lactone (iPLA2 inhibitor); 0.1 mM for Valeroyl salicylate (PTGS1 inhibitor) and 4 μM for Celecoxib (PTGS2 inhibitor) and prostaglandins PGD2, PGF2α, and PGE2 all used at 0.022 µM. **b)** MTT assay was used to test the toxicity of the PTGER4 agonist and antagonist in mouse organoids. The treatments were: Control, PTGER4 agonist L-902,688 (902) at 1 micromolar and 10 micromolar; and the antagonist L- 161,982 (161) at 10 micromolar and at 20 micromolar. Experiments were carried out using 2 mouse colons for generate the organoids. Values represent mean ± SEM.

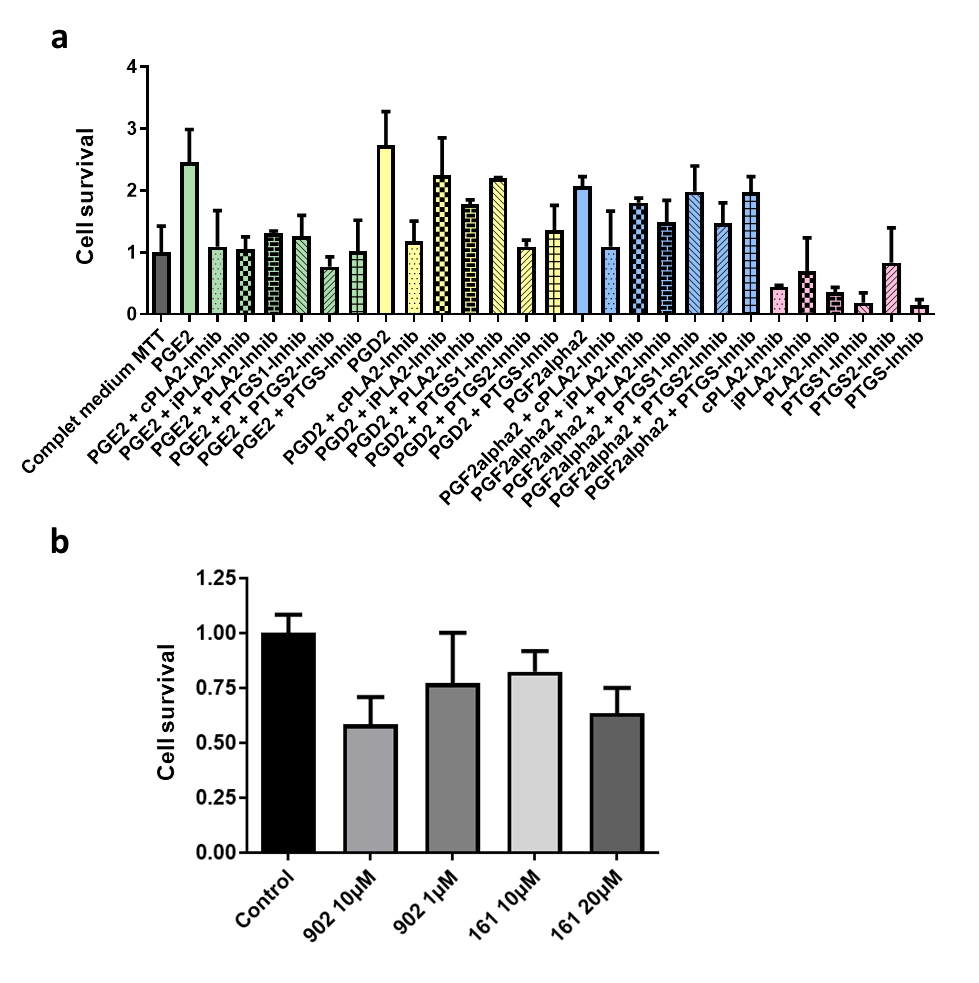

#### **Supplementary Figure 10**. Effect on organoid growth rate at 48h upon PTGER4 agonist and antagonist, PTGS and PLA2 inhibition and prostaglandin treatments. 5 μM for Arachidonyl trifluoromethyl ketone (cPLA2 inhibitor); 20 μM for Bromoenol lactone (iPLA2 inhibitor); 0.1 mM for Valeroyl salicylate (PTGS1 inhibitor) and 4 μM for Celecoxib (PTGS2 inhibitor), 1 µM for L-902,688 and 10 µM L-161.982. PGD2, PGF2α, and PGE2 all used at 0.022 µM.

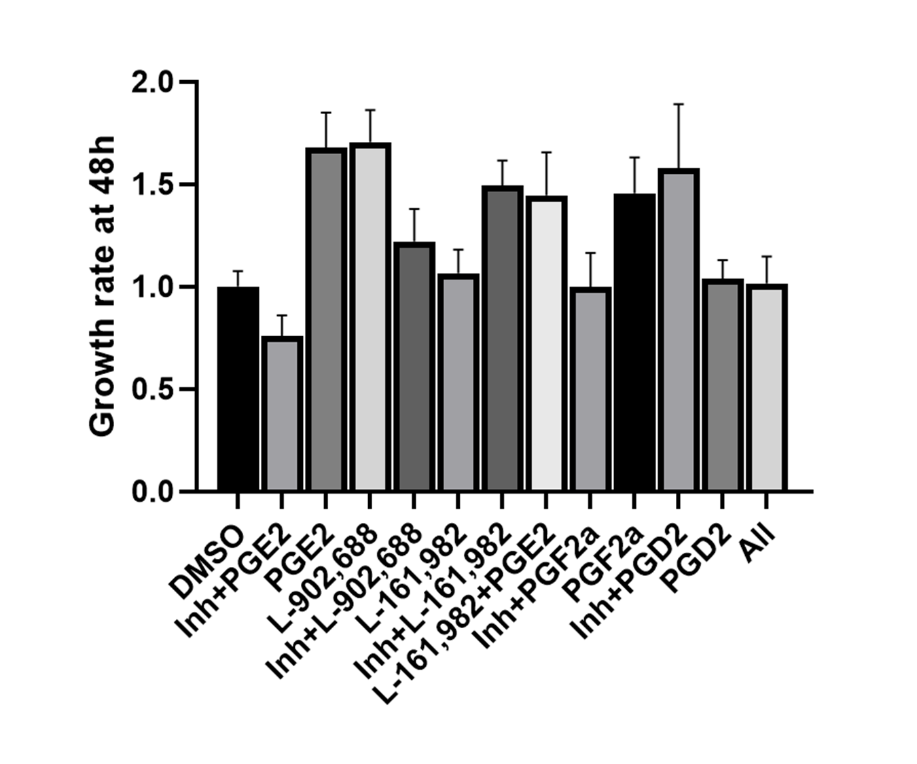

#### **Supplementary Figure 11.** Tuft cell markers DCLK1, HPGDS, PTGS1, and KIT gene expression decreases in colon cancer samples compared to normal tissue. Cohort: TCGA Colon and Rectal Cancer (COADREAD), dataset: HiSeqV2, Consensus molecular subtype (CMS) samples = 349, gene expression RNAseq data, unit log2(norm_count+1) CMS labels were obtained from Guinney et al., 2015. Data was analyzed and plotted with xenabrowser.net^a^.

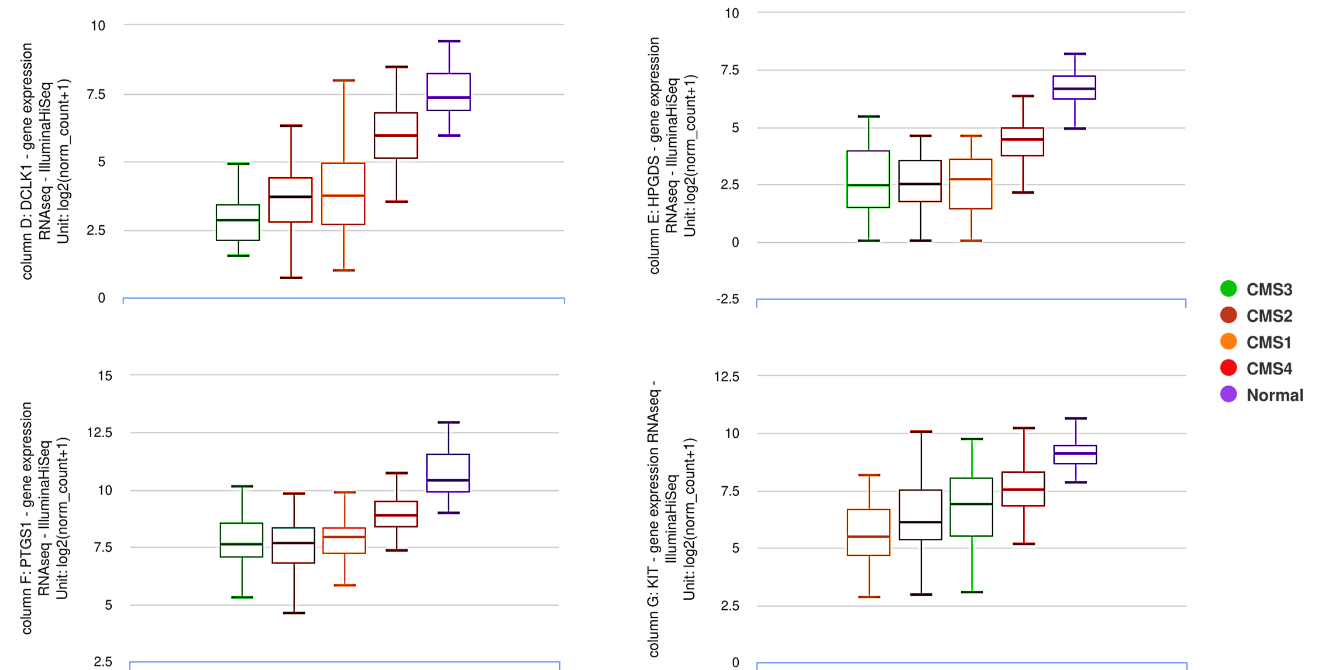

^a^ Goldman, M.J. et al. (2020). Visualizing and interpreting cancer genomics data via the Xena platform. Nat. Biotechnol. *38*, 675–678. https://doi.org/10.1038/s41587-020-0546-8

#### **Supplementary Figure 12.** Representative configuration of FACSAria Fusion with FACSDiva v8.0 (BD Biosciences) gating for EPHB2 subpopulations. Healthy (a) and tumor (b) surgical samples of colon cancer patient 321. The procedure was followed as previously described. Briefly, EpCAM positive (CD31, -45, -11b, and -117 negative cells were sorted based on EPHB2 intensity in 4 groups (Negative, Low, Medium, and High). Samples were collected into 300 μl of PBS for MALDI-IMS, and 300 μl of RLT buffer (QIAGEN) for gene expression microarray.

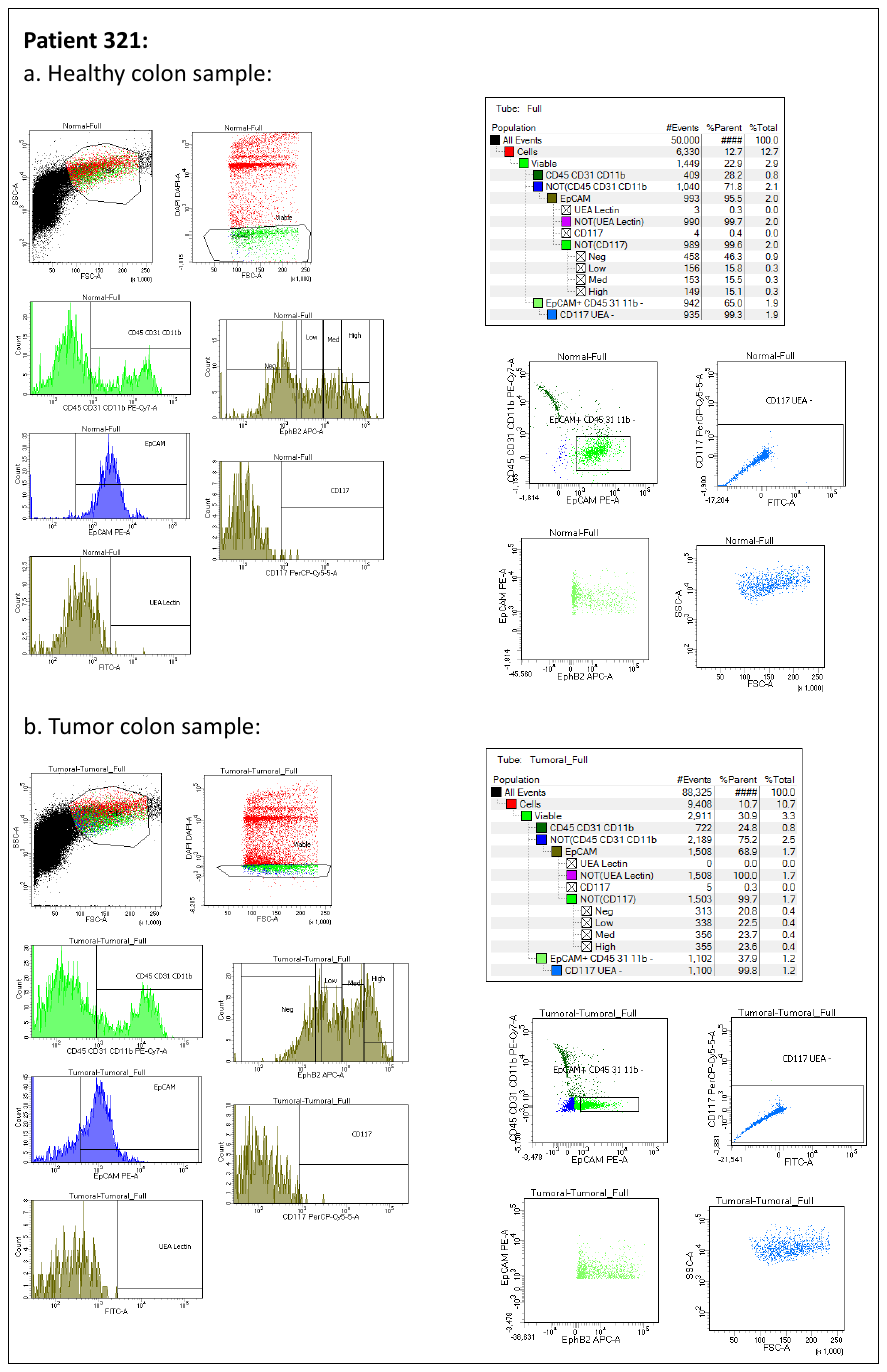

#### **Supplementary Figure 13.** Weighted Gene Co-expression Network Analysis**:** a soft-thresholding power was selected to ensure the scale free topology, with a β-value of 12 (scale-free R^2^ = 0.90) for the healthy samples; and a β-value of 16 (scale-free R^2^ = 0.50 - 0.60) for the tumor. Minimum-module-size was set to 30 probes, and modules whose distance is less than 0.25 were merged.

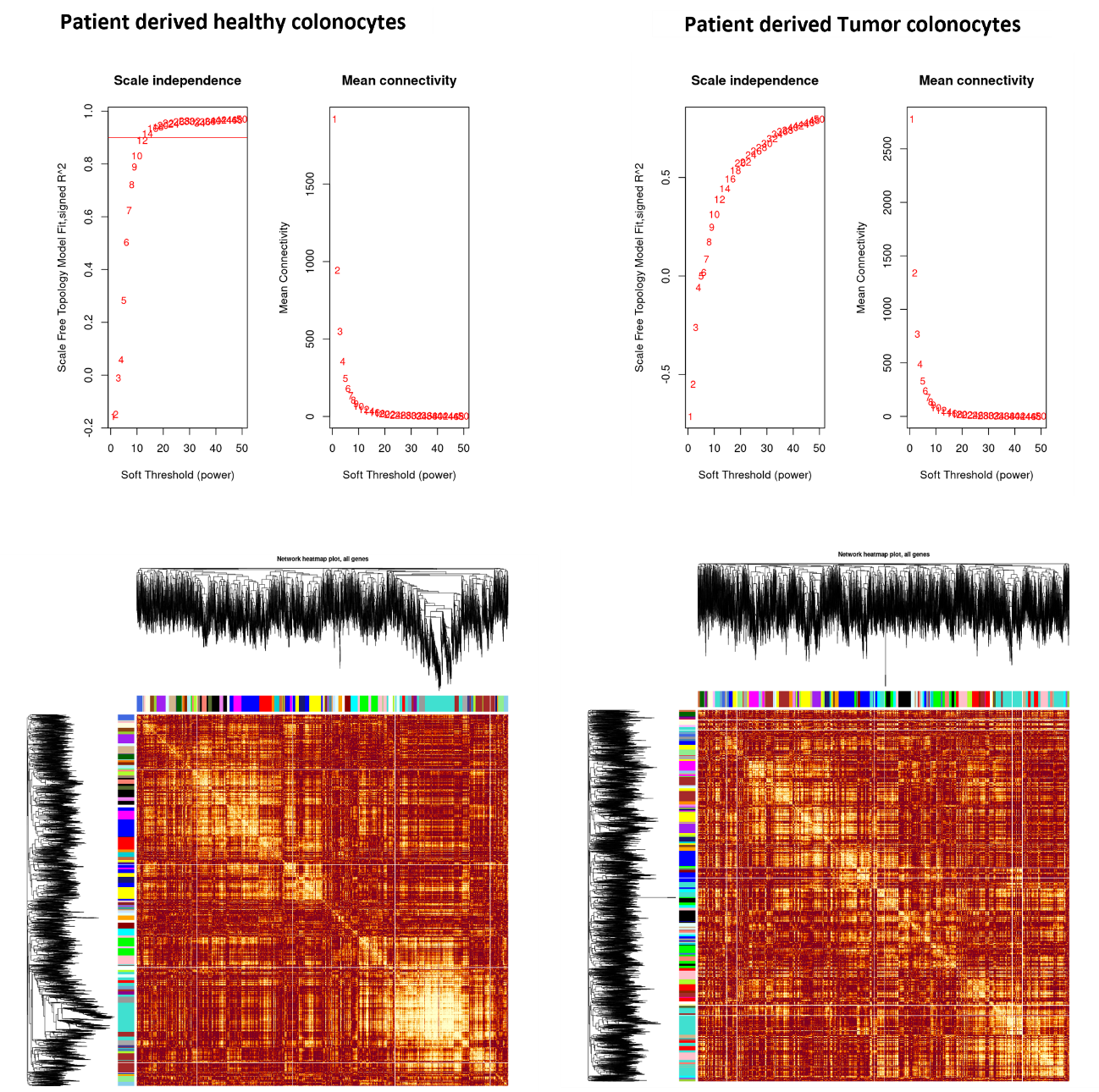
